# Lipid flippase dysfunction as a novel therapeutic target for endosomal anomalies in Alzheimer’s disease

**DOI:** 10.1101/2021.07.30.454423

**Authors:** Nanaka Kaneshiro, Masato Komai, Ryosuke Imaoka, Atsuya Ikeda, Yuji Kamikubo, Tadafumi Hashimoto, Takeshi Iwatsubo, Takashi Sakurai, Takashi Uehara, Nobumasa Takasugi

## Abstract

β-amyloid precursor protein (APP) and their metabolites are deeply involved in the development of Alzheimer’s disease (AD). Upon the upregulation of β-site APP cleaving enzyme 1 (BACE1), its product, the β-carboxyl-terminal fragment of APP (βCTF), is accumulated in the early stage of sporadic AD brains. βCTF accumulation is currently considered the trigger for endosomal anomalies to form enlarged endosomes, one of the earliest pathologies in AD. However, the details of the underlying mechanism remain largely unclear. In this study, using BACE1 stably-overexpressing cells, we describe that lipid flippase subcomponent TMEM30A interacts with accumulated βCTF. Among the lipid flippases in endosomes, those composed of TMEM30A and active subcomponent ATP8A1 transports phospholipid, phosphatidylserine (PS), to the cytosolic side of the endosomes. The lipid flippase activity and cytosolic PS distribution are critical for membrane fission and vesicle transport. Intriguingly, accumulated βCTF in model cells impaired lipid flippase physiological formation and activity, along with endosome enlargement. Moreover, in the brains of AD model mice before the amyloid-β (Aβ) deposition, the TMEM30A/βCTF complex formation occurred, followed by lipid flippase dysfunction. Importantly, our novel Aβ/βCTF interacting TMEM30A-derived peptide “T-RAP” improved endosome enlargement and reduced βCTF levels. These T-RAP effects could result from the recovery of lipid flippase activity. Therefore, we propose lipid flippase dysfunction as a key pathogenic event and a novel therapeutic target for AD.

## Introduction

Amyloid-β (Aβ) peptides are accumulated in the brains of patients with Alzheimer’s disease (AD)^1^ and are produced from the sequential cleavage of the β-amyloid precursor protein (APP) by the β-site APP cleaving enzyme 1 (BACE1)^2,3,4^ and γ-secretase^5^. Although pieces of genetic and biological evidence suggest the link between Aβ and AD pathogenesis, several clinical trials based on the “amyloid hypothesis” have failed^6^. Therefore, new ideas and therapeutic targets that complement the hypothesis are required for this field.

A recent report identified that traffic impairment, showing endosomal anomalies, is an early pathogenic event in AD before Aβ deposition^7^. In line with this, several studies in AD model mice^8^, human iPSC-derived neurons^9^, and AD brains^10^ have shown that the accumulated β-carboxyl-terminal fragment of APP (βCTF), the product of BACE1 and direct precursor of Aβ, is the cause of endosomal anomalies. Indeed, BACE1 expression and activity are upregulated at the early stage of sporadic AD^11,12^. However, the details of the mechanism underlying the βCTF mediated endosomal anomalies remain unclear.

Previously, we identified TMEM30A (CDC50A), a subcomponent of lipid flippases, as a candidate partner for βCTF. This complex mediates the formation of enlarged endosomes^13^. Most lipid flippases consist of TMEM30A and active subcomponents, P4-ATPases. These enzymes translocate phospholipids from the exoplasmic/luminal side to the cytoplasmic leaflet of the lipid bilayer to regulate phospholipids asymmetry^14^.

Phosphatidylserine (PS), one of the phospholipids, is a component of lipid bilayer and mainly localizes at the cytoplasmic leaflet^14^. PS level on the cytosolic side in endosomes determines the recruitment of PS-binding proteins to trigger membrane budding, which promotes subsequent vesicle fission and transport^15,16^. This PS asymmetry is regulated by one of the endosomal lipid flippases, those composed of TMEM30A and active subcomponent ATP8A1, which has a high affinity for PS^14^. These lines of evidence indicate that lipid flippase activity is essential for vesicular trafficking.

In this study, we investigated whether lipid flippase activity contributes to the βCTF-mediated endosomal anomalies and could be a novel therapeutic target for AD treatment.

## Results

### BACE1 upregulation promotes complex formation between endogenous TMEM30A and accumulated βCTF, and endosomal anomalies

To gain insight into the early pathology of AD, we established BACE1 stably-overexpressing SH-SY5Y cell lines (SH-BACE1 cells). BACE1 produces β1 or β11CTF depending on APP cleavage sites^17^. BACE1 upregulation increased the β-secretase products sAPPβ, βCTF, and Aβ, although it severely reduced α-secretase-cleaved carboxyl-terminal fragment (αCTF), in good agreement with previous reports^18^ (Fig. 1A and Supplementary Fig. 1A). We previously demonstrated that TMEM30A is a candidate partner for βCTF (β1/β11CTF)-mediated endosomal anomalies^13^. Supportively, CFP-TMEM30A, SC100^19^, and SC89 (artificial β1 and β11CTF) co-transfection revealed that TMEM30A could interact with these βCTF (Supplementary Fig. 1B). Although βCTF in non-treated SH-BACE1 cells might deserve further analysis, we found that endogenous βCTF exhibited resistance to mild detergents such as CHAPS (Supplementary Fig. 1C and D). Indeed, βCTF insolubility in mild detergents was reported elsewhere^20^. Our result suggests that the environment surrounding βCTF is particularly unique. Therefore, to verify whether endogenous TMEM30A interacts with accumulated βCTF in SH-BACE1 cells, we treated γ-secretase inhibitor DAPT to accumulate APP-CTF, then performed co-immunoprecipitation analysis. In line with our previous findings, we observed an interaction between the endogenous TMEM30A and accumulated βCTF, but not αCTF, and the BACE1 inhibitor co-treatment abolished this interaction (Fig. 1B).

**Figure 1.**
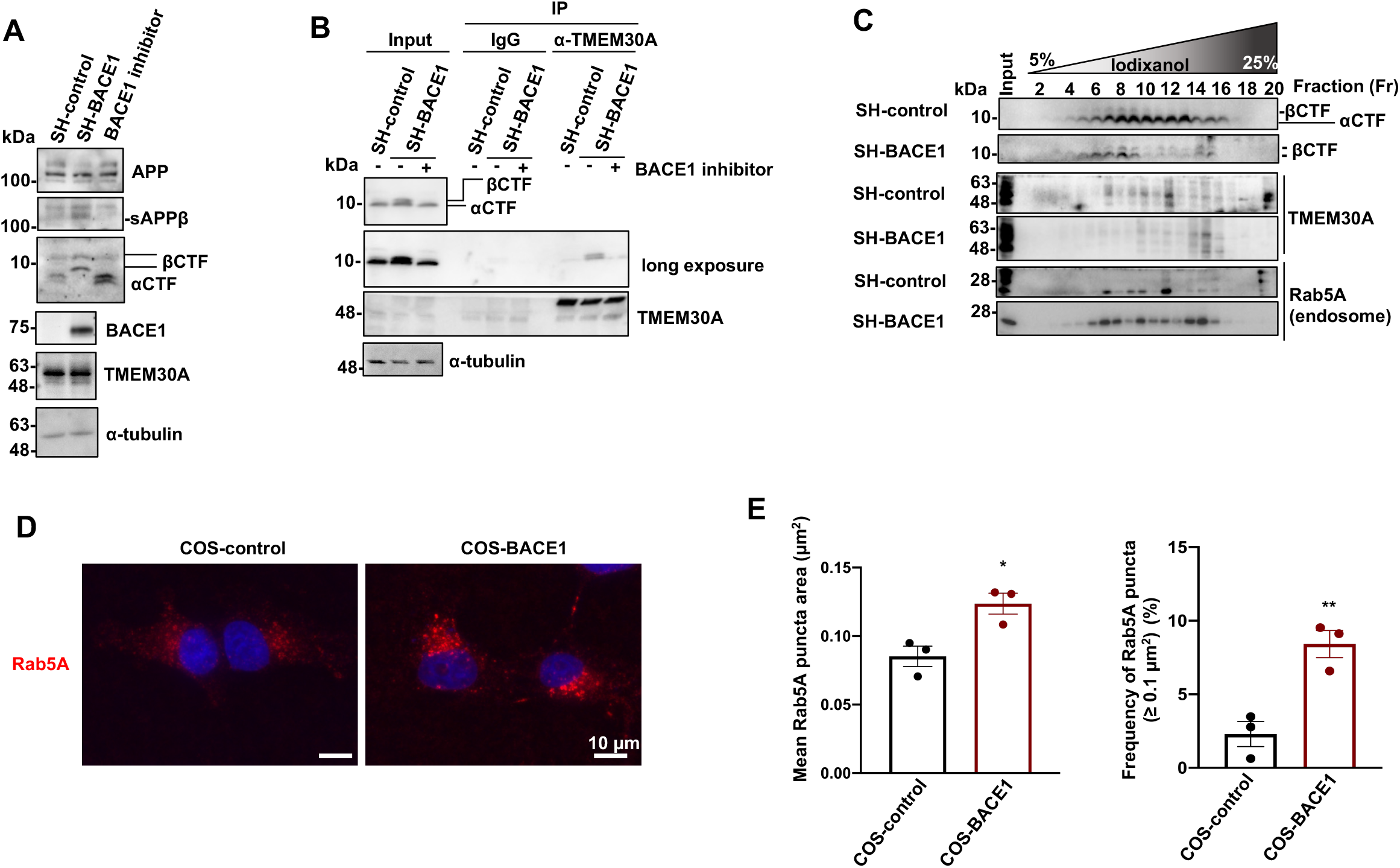
BACE1 upregulation induces the complex formation between TMEM30A and βCTF and endosomal anomalies. (**A**) Immunoblotting analysis for APP metabolites and TMEM30A in SH-control and SH-BACE1 cells. β-secretase inhibitor IV (10 µM) was treated to SH-BACE1 cells. (**B**) Co-immunoprecipitation analysis using TMEM30A antibody in the accumulation of βCTF by the DAPT (10 µM) treatment or co-treatment with β-secretase inhibitor IV (10 µM) for 24 h. (**C**) Iodixanol gradient fractionation of the homogenates from SH-control and SH-BACE1 cells. For the endosome marker, Rab5A was used. (**D**) COS-control and COS-BACE1 cells were immunostained for Rab5A (Red). DAPI stained the nucleus (Blue). Scale bars: 10 µm. Representative z-stack images were captured using a 60x objective lens (zoom x2.6). (**E**) Quantitation of size distribution (≥ 0.1 μm^2^) and mean area of Rab5A-positive puncta (*n*=3, mean ± SEM, two-tailed Student’s t-test, \**P*<0.05, *\*\*P*<0.01).

Next, a stepwise iodixanol gradient organelle fractionation was performed as described previously^21^ to analyze the protein distributions. The distribution of LAMP2 (lysosome marker) and calreticulin (ER marker) showed no obvious change between SH-control and SH-BACE1 cells (Supplementary Fig. 2A). In contrast, Rab5A, an early endosome marker, was distributed in fractions (Fr) 7∼12 in SH-control cells (Fig. 1C) and its distribution was broadened to the heavier fractions (Fr 14∼16) in SH-BACE1 cells. Similarly, βCTF, enriched in Fr 6∼9 in SH-control cells, changed its distribution in the heavier fractions (Fr 14∼16) in SH-BACE1 cells (Fig. 1C). Concomitantly, BACE1 was distributed in Fr 14∼16 in SH-BACE1 cells (Supplementary Fig. 2A). Intriguingly, TMEM30A was mainly distributed in Fr 7∼12 in SH-control cells. However, in SH-BACE1 cells, its distribution drastically changed to Fr 14∼16 where Rab5, βCTF, and BACE1 abnormally co-localized (Fig. 1C). We observed no alteration in the TMEM30A and Rab5A protein levels between SH-control and SH-BACE1 cells (Fig. 1A and Supplementary Fig. 2B).

Next, we performed immunofluorescence analysis using BACE1 stably-overexpressing COS-7 cells (COS-BACE1), well-characterized in organelle morphology observations^13,22,23^. In contrast to the lack of alteration in the Rab5A protein level (Supplementary Fig. 2C), the mean of the Rab5A-positive puncta area and the frequency of size distribution (≥ 0.1 μm^2^) were significantly increased in COS-BACE1 cells (Fig. 1D, E, and Supplementary Fig. 2D).

Our findings suggest that BACE1 increases βCTF level, which interacts with TMEM30A and mediates endosomal anomalies.

### βCTF accumulation triggers the lipid flippase activity impairment

TMEM30A interacts with active subunits, P4-ATPases, to form lipid flippase, and contributes to stability, distribution, and activity of P4-ATPases^14,24^. Therefore, we hypothesized that lipid flippase activity is associated with TMEM30A/βCTF-mediated endosomal anomalies.

Immunoprecipitation analysis in the membrane fractions revealed that the physiological complex formation between TMEM30A and ATP8A1, an endosomal P4-ATPase which has a high affinity for PS, significantly decreased in SH-BACE1 cells. The BACE1 inhibitor treatment recovered this interaction in SH-BACE1 cells (Fig. 2A, B, and Supplementary Fig. 3A), indicating that this deficit is BACE1 activity-dependent.

**Figure 2.**
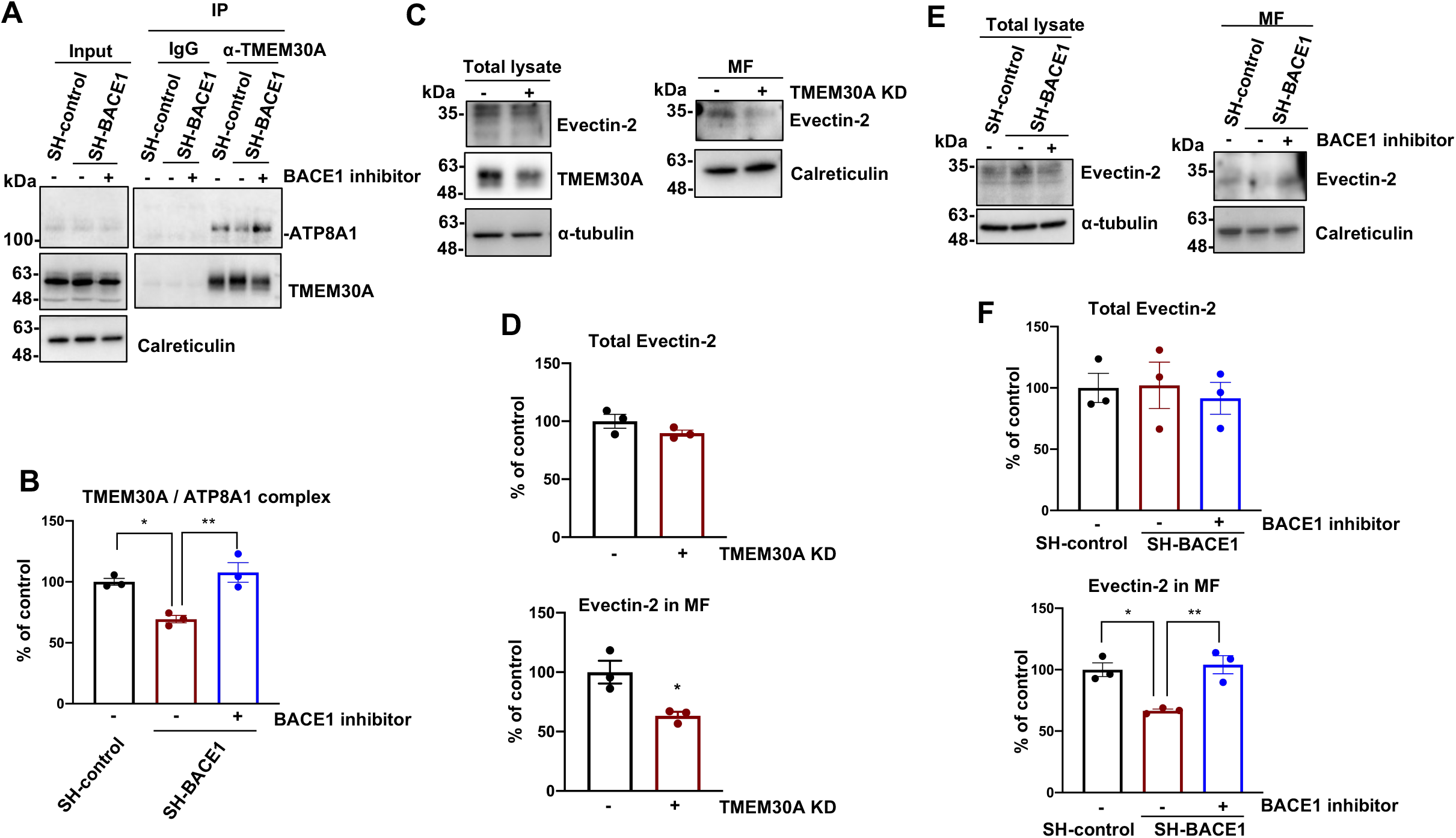
BACE1 upregulation induces lipid flippase dysfunction depending on the BACE1 activity. (**A**) The membrane fractions from SH-control and SH-BACE1 cells applied to co-immunoprecipitation analysis using TMEM30A antibody. Cells were treated with the β-secretase inhibitor IV (10 µM) for 48 h. Calreticulin was used as a loading control of membrane fractions. (**B**) Quantification of ATP8A1 co-immunoprecipitated by TMEM30A antibody in Fig. 2A (*n*=3, mean ± SEM, one-way ANOVA Bonferroni’s multiple comparisons test, \**P*<0.05, \*\**P*<0.01). (**C**) Immunoblotting analysis for Evectin-2 in total cell lysates and membrane fractions (MF) in the knockdown of TMEM30A in SH-SY5Y cells. (**D**) Quantification of Evectin-2 localization in whole cell lysates or MF (*n*=3, mean ± SEM, two-tailed Student’s t-test, \**P*<0.05). (**E**) Immunoblotting analysis for Evectin-2 in total cell lysates and MF in the treatment of the β-secretase inhibitor IV (10 µM) for 48 h. (**F**) Quantification of the Evectin-2 localization in total lysates or MF (*n*=3, mean ± SEM, one-way ANOVA Bonferroni’s multiple comparisons test, \**P*<0.05, ** *P*<0.01).

Since it is difficult to evaluate lipid flippase activity in organelles such as endosomes, we attempted to exploit the endosomal PS binding property of Evectin2, a promoting factor of membrane fission in vesicle transport to analyze the lipid flippase activity. Evectin-2 has a PS-specific pleckstrin-homology (PH) lipid-binding domain, and its association with the endosomal membrane depends on lipid flippase activity^23,25^. As previously reported^23^, the predominant distribution of Evectin-2 in the membrane fractions was abolished by TMEM30A knockdown (Fig. 2C, D, and Supplementary Fig. 3B). Intriguingly, Evectin-2 dissociated from the membrane fractions in SH-BACE1 cells and got redistributed upon the BACE1 inhibitor treatment without altering total Evectin-2 level (Fig. 2E and F).

To further quantify Evectin-2 localization in the endosomes, we used a NanoBiT luciferase reconstitution system. Large BiT (LgBiT) and Small BiT (SmBiT) are the parts of the *Oplophorus gracilirostris-*derived Nanoluc luciferase. As SmBiT has a low affinity (Kd = 190 μM) to the LgBiT, their interaction is fragile and reversible, and the luciferase activity is not reconstituted without their forced proximity by fused proteins^26^. We fused these subunits to Rab5A (LgBiT-Rab5) and Evectin-2 (SmBiT-Evectin-2) to monitor the Evectin-2 localization in the endosomes (Fig. 3A). Immunofluorescence analysis showed that LgBiT-Rab5 localized in the endosomes (Supplementary Fig. 4A). Moreover, similar to our biochemical analysis (Fig. 2C and D), TMEM30A knockdown significantly decreased the luciferase activity (Fig. 3B, C and Supplementary Fig. 4B). For further validation, we investigated the effect of Evectin-2 mutant on the PH domain (K20E), lacking the binding affinity to PS^23^. We observed significantly reduced luciferase activity in the K20E mutant (Supplementary Fig. 4C and D). These results clearly show that our reporter system could be used for semi-quantitative estimation of lipid flippase activity in endosomes.

**Figure 3.**
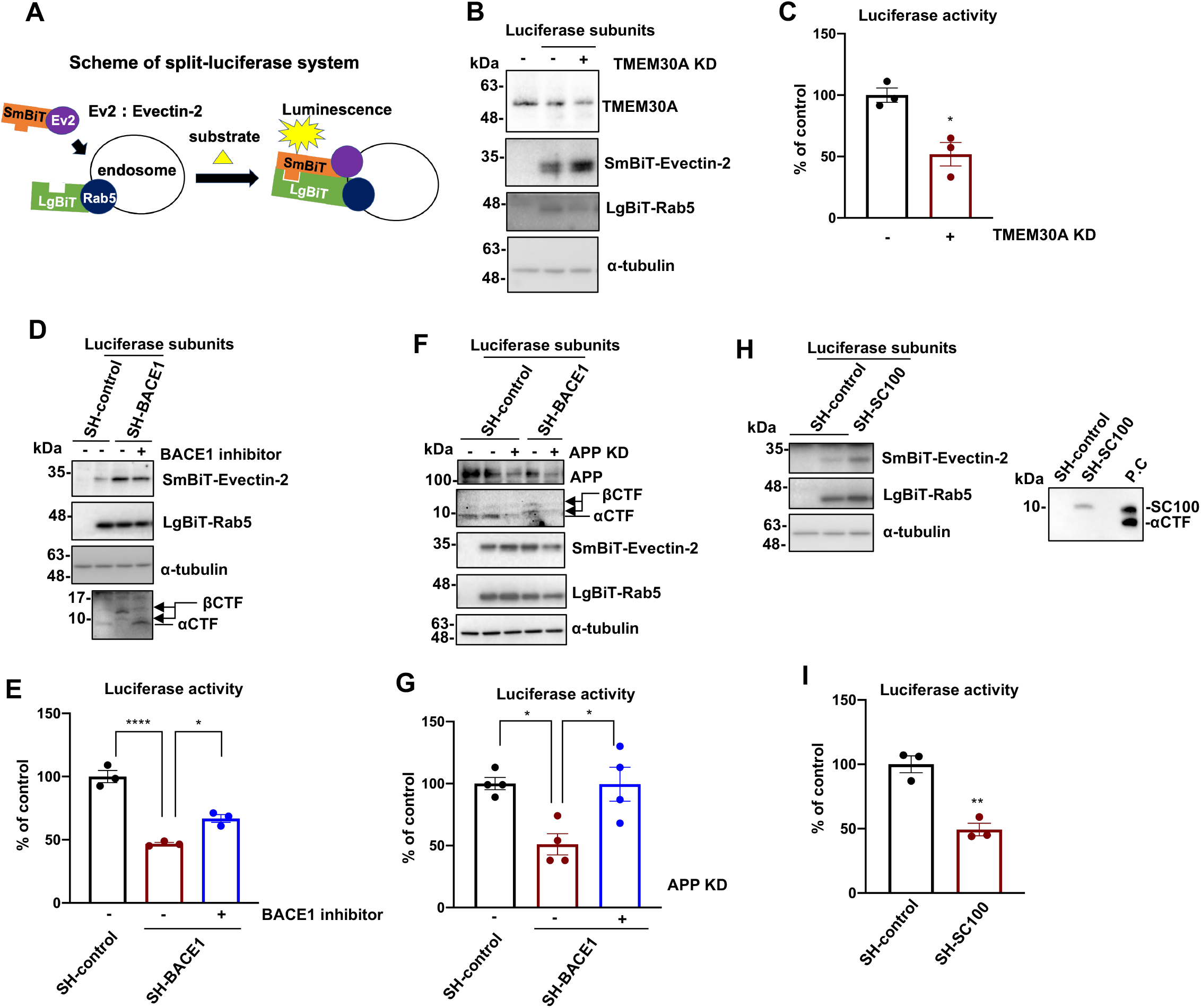
The semi-quantitative analysis shows the lipid flippase activity in endosomes is decreased depending on the levels of βCTFs. (**A**) Schematic view of the measurement methods for endosomal lipid flippase activity using a split-luciferase assay system. Luciferase subunits, LgBiT or SmBiT, were fused with Rab5A or Evectin-2, respectively. The reconstitution of luciferase activity depends on the endosomal distribution of Evectin-2, which reflects the endosomal lipid flippase activity (Fig.2). (**B**) Immunoblotting analysis for LgBiT-Rab5, SmBiT-Evectin-2, and TMEM30A using HA, FLAG, and TMEM30A antibodies in the knockdown of TMEM30A for 72 h in SH-SY5Y cells. (**C**) Quantification of the luciferase activity in the knockdown of TMEM30A for 72 h in SH-SY5Y cells (*n*=3, mean ± SEM, two-tailed Student’s t-test, \**P*<0.05). (**D**) Immunoblotting analysis for LgBiT-Rab5, SmBiT-Evectin-2, and APP-CTF in the treatment of β-secretase inhibitor IV (10 µM) for 48 h. (**E**) Quantification of the luciferase activity for treating β-secretase inhibitor IV (10 µM) for 48 h (*n*=3, mean ± SEM, one-way ANOVA Bonferroni’s multiple comparisons test, \**P*<0.05, \*\*\*\**P*<0.0001). (**F**) Immunoblotting analysis for LgBiT-Rab5, SmBiT-Evectin-2, APP, and APP-CTF in the knockdown of APP for 72 h. (**G**) Quantification of the luciferase activity in the knockdown of APP for 72 h (*n*=4, mean ± SEM, one-way ANOVA Bonferroni’s multiple comparisons test, \**P*<0.05). (**H**) Immunoblotting analysis for SC100, LgBiT-Rab5, and SmBiT-Evetin-2. P.C is the control for SC100 and αCTF. (**I**) Quantification of the luciferase activity in SH-control and SH-SC100 cells 48 h after the transfection of luciferase subunits (*n*=3, mean ± SEM, two-tailed Student’s t-test, \*\**P*<0.01).

Next, we applied this system to SH-BACE1 cells. Intriguingly, we observed a significant reduction in the lipid flippase activity, rescued by the BACE1 inhibitor treatment (Fig. 3D and E). To confirm that the BACE1-mediated APP cleavage is a prerequisite for this event, we performed APP knockdown in SH-BACE1 cells and detected lipid flippase activity recovery (Fig. 3F, G and Supplementary Fig. 5A-C). Interestingly, the APP knockdown failed to affect the lipid flippase activity in SH-control cells (Supplementary Fig. 5D). Furthermore, lipid flippase activity significantly decreased in SC100 stably overexpressing SH-SY5Y cells (Fig. 3H and I).

Our data strongly support the hypothesis that βCTF accumulation leads to lipid flippase dysfunction via abnormal complex formation with TMEM30A.

### TMEM30A/βCTF complex formation and subsequent lipid flippase dysfunction precede Aβ deposition in AD model mice

Next, we explored the TMEM30A/βCTF complex formation and lipid flippase dysfunction at the early stage in AD model mice. As A7 model mice show a relatively slow Aβ deposition, starting at approximately 9 months of age, we used these AD model mice as an appropriate model to observe the precursory phenomenon^27^. At 3 and 6 months, the ATP8A1, TMEM30A, and Evectin-2 protein levels were not significantly different between the WT and transgenic (Tg) mice (Supplementary Fig. 6A and B). Moreover, we observed no significant difference in the αCTF and βCTF levels between the 3- and 6-month-old Tg mice (Supplementary Fig. 6A and B). Intriguingly, TMEM30A interacted with βCTF in both 3- and 6-month-old model mice (Fig. 6A and B). However, TMEM30A failed to interact with Aβ oligomers, previously described as another factor in vesicular traffic impairment^28^, and with Aβ monomers (Supplementary Fig. 7). Importantly, both lipid flippase formation (Fig. 4A and C) and Evectin-2 localization in the membrane fractions (Fig. 4A and D) significantly decreased in 6-month-old A7 mice. We confirmed that the indicated band is Evectin-2 (Supplementary Fig. 6C). Our results suggest that the TMEM30A/βCTF complex induces lipid flippase dysfunction, which precedes Aβ deposition.

**Figure 4.**
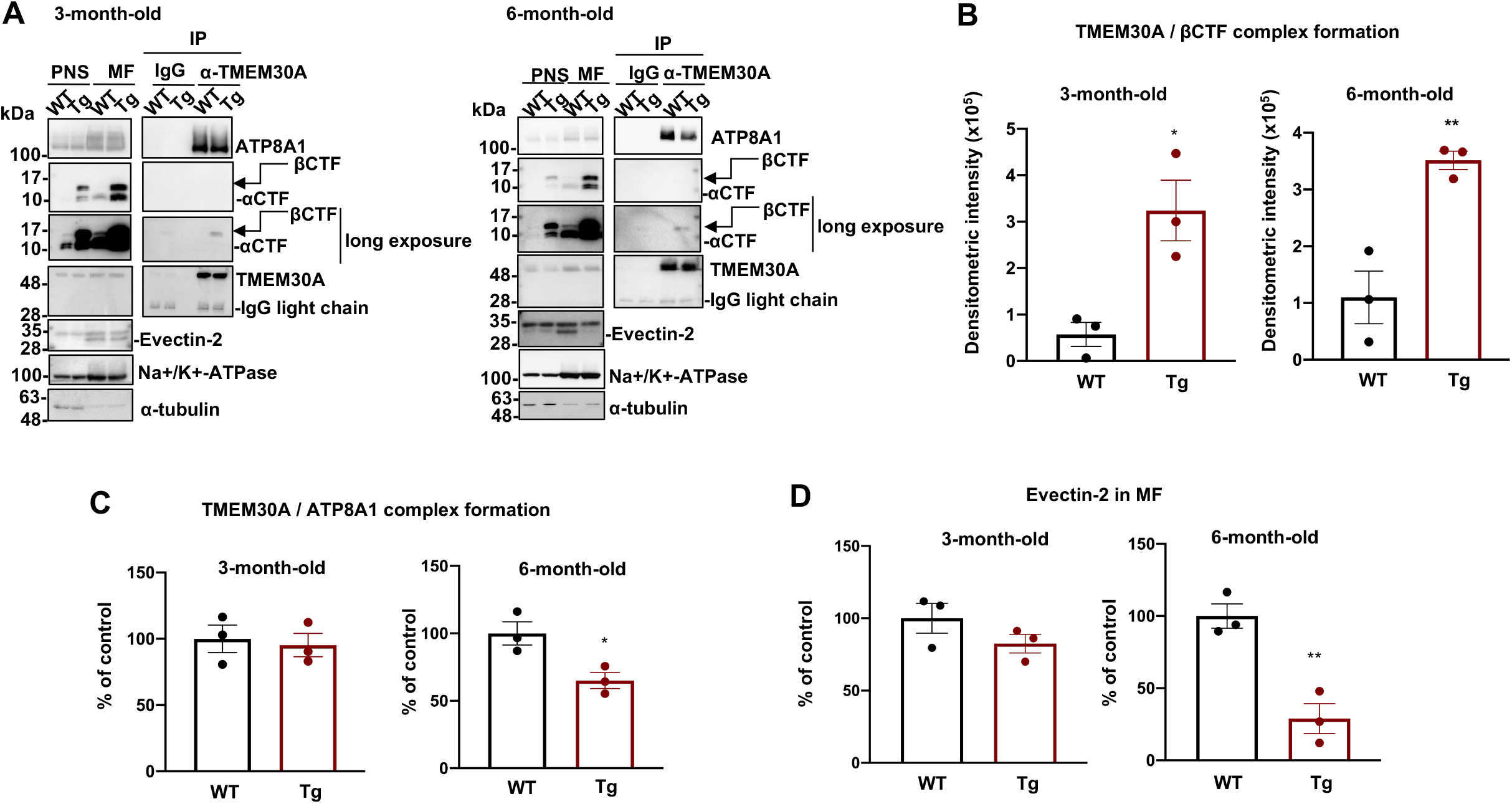
TMEM30A interacts with βCTF in A7 mice, which follows by lipid flippase dysfunction. (**A**) The membrane fractions from WT and A7 mice brain at 3- or 6-month-old applied to co-immunoprecipitation analysis using TMEM30A antibody. Na+/K+-ATPase was used as a loading control of membrane fractions. (**B, C**) Quantification of the complex formation of (**B**) TMEM30A and βCTF or (**C**) TMEM30A and ATP8A1 (*n*=3, mean ± SEM, two-tailed Student’s t-test, \**P*<0.05, \*\**P*<0.01). (**D**) Quantification of Evectin-2 localization in membrane fractions (*n*=3, mean ± SEM, two-tailed Student’s t-test, \*\**P*<0.01).

### βCTF/Aβ interactive peptide “T-RAP” improves endosome enlargement

We expected that the inhibition of the interaction between TMEM30A and βCTF could improve lipid flippase dysfunction and endosomal anomalies. Previously, we identified that the extracellular domain of TMEM30A (TmEx) interacts with the Aβ sequence of βCTF^13^. We explored the interacting domain by sequential deletion of TmEx using GST-pulldown assay (Fig. 5A-C) and found that the 117–166 AA region contains the critical residues for the interaction with βCTF (Fig. 5B-D). Using RaptorX prediction (http://raptorx.uchicago.edu/), we noticed that the conformation of the α-helices and β-sheet exists in the well-conserved 125–150 AA region. We named this sequence “T-RAP” (<u>T</u>mem30A <u>r</u>elated <u>a</u>myloid-beta interacting <u>p</u>eptide) (Fig. 5A). We verified that GST-fused T-RAP efficiently pulled down βCTF and Aβ, but not αCTF (Fig. 5B, D and Supplementary Fig. 8), indicating T-RAP has a high affinity for Aβ N-terminal sequence.

**Figure 5.**
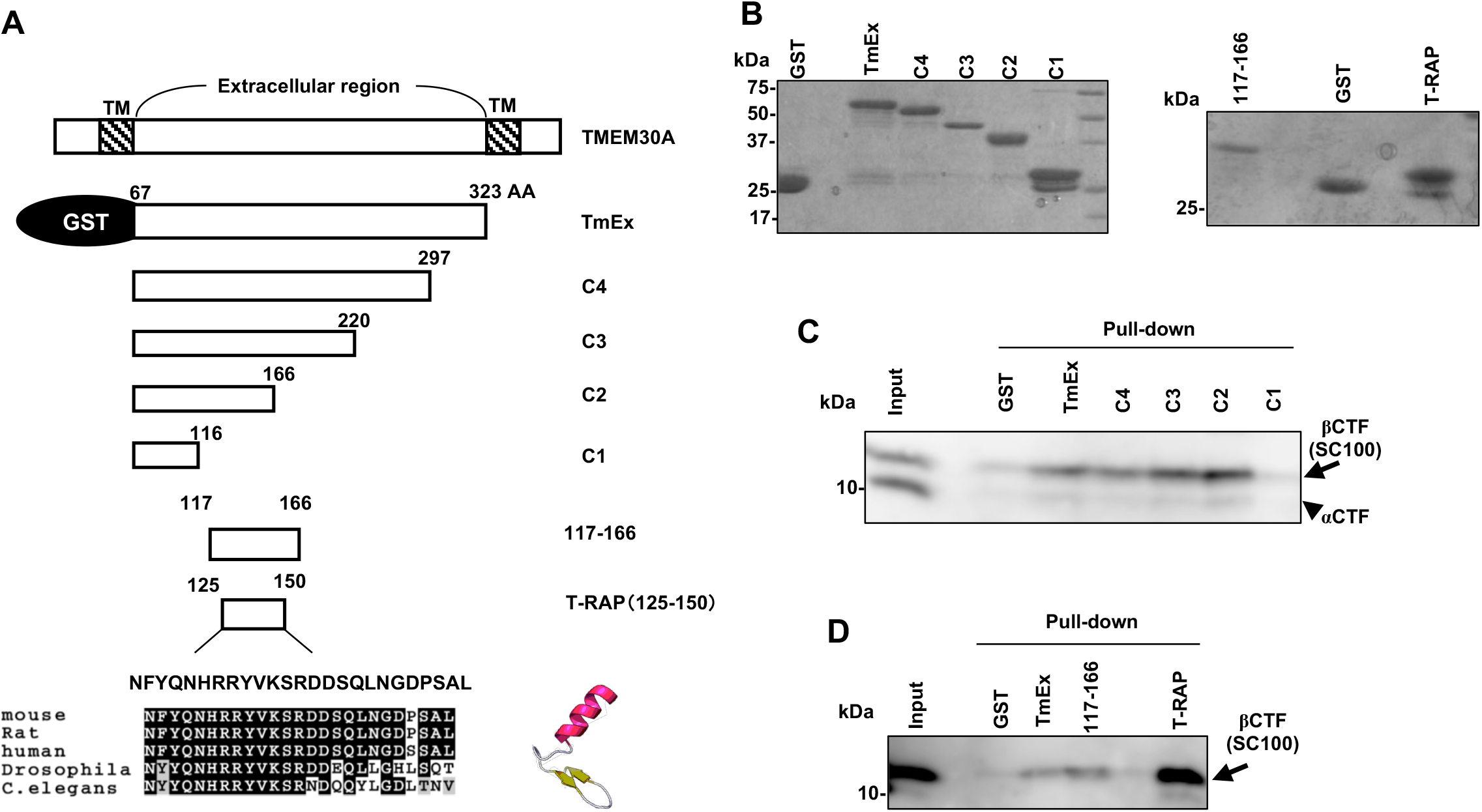
Identification of T-RAP peptide. (**A**) Schematic view of the sequential deletion constructs of GST-TmEx (Extracellular-domain of TMEM30A) and GST fused TMEM30A (117-166 AA) and “T-RAP” sequence. Lowest panel: The conservation of the T-RAP sequence among indicated species and predicted structure by Raptor-X. (**B**) Coomassie’s brilliant blue staining of purified GST-fusion proteins in Fig 5A. (**C, D**) GST-pull down from HEK293 lysate transfected with SC100.

To assess whether the synthetic T-RAP peptide influences lipid flippase activity, we treated SH-SY5Y cells with T-RAP and performed a flippase activity assay. In advance, we confirmed that T-RAP displayed no toxicity at the density used in this study (Supplementary Fig. 9A). Although not significant, T-RAP showed a trend to improve lipid flippase activity in SH-BACE1 cells (Fig. 6A and B). On the other hand, T-RAP (50 μM) fused with the Trans-Activator of Transcription Protein (TAT) which is easily introduced into cells significantly rescued the lipid flippase activity in SH-BACE1 cells (Supplementary Fig. 9B). We consider that TAT-T-RAP could penetrate membranes more efficiently than T-RAP.

**Figure 6.**
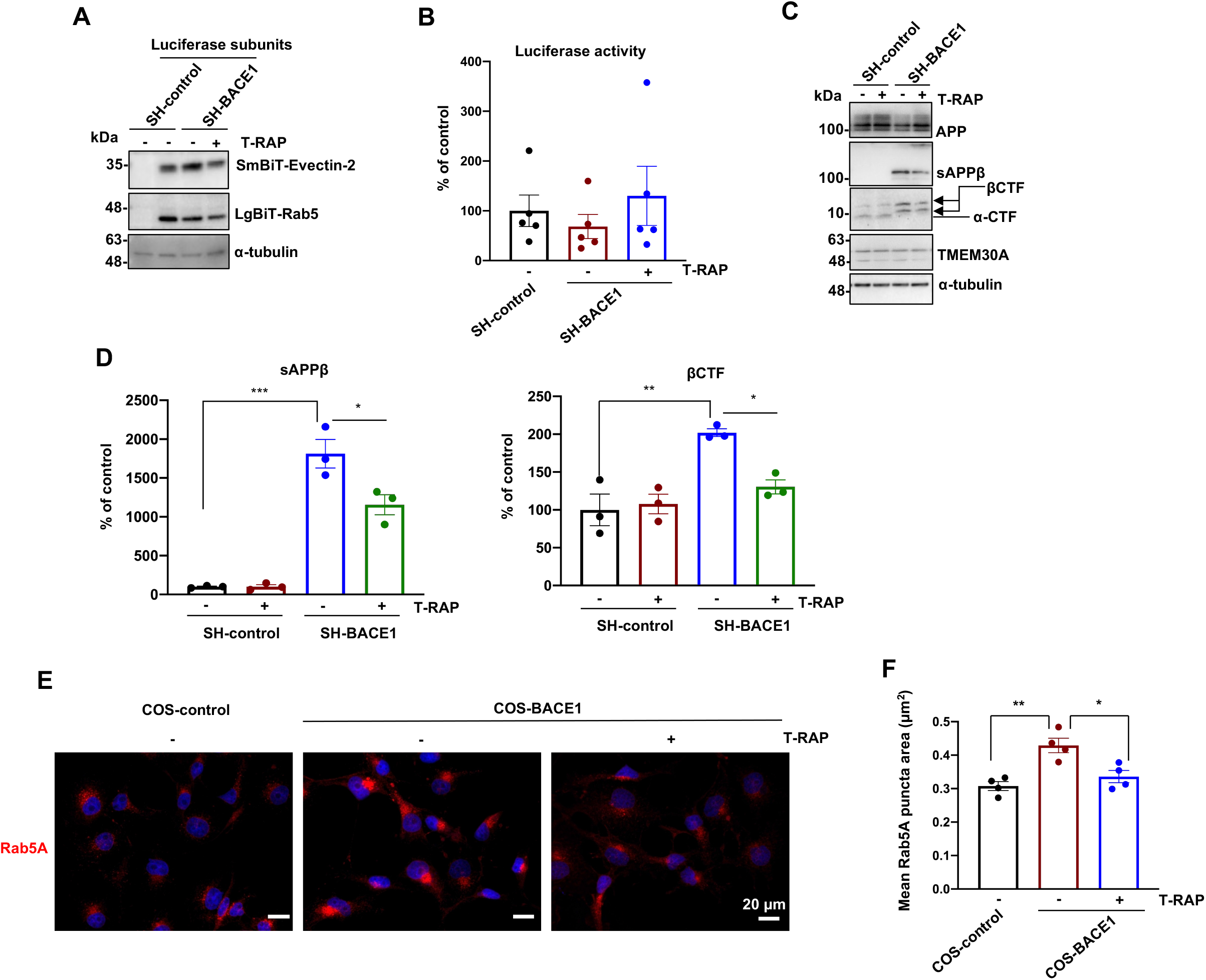
βCTF interacting peptide ‘T-RAP’ shows a trend to rescue lipid flippase activity and improves the endosomal anomalies in BACE1 upregulation. (**A**) Immunoblotting analysis for SmBiT-Evectin-2 and LgBiT-Rab5 after the treatment of T-RAP (10 µM) for 48h. (**B**) Quantification of the luciferase activity in T-RAP (10 µM) treatment for 48 h (*n*=5, mean ± SEM, one-way ANOVA Bonferroni’s multiple comparisons test). (**C**) Immunoblotting analysis for APP metabolites and TMEM30A in T-RAP (10 µM) treatment for 48 h. (**D**) Quantification of sAPPβ and βCTF in Fig. 6C (*n*=3, mean ± SEM, one-way ANOVA Bonferroni’s multiple comparisons test, \**P*<0.05, \*\**P*<0.01, \*\*\**P*<0.001). (**E**) COS-control and COS-BACE1 cells were immunostained for Rab5A after T-RAP (10 µM) treatment for 48 h. Scale bars: 20 µm. Representative z-stack images were captured using a 60x objective lens. (**F**) Quantification of the mean Rab5A positive puncta area (*n*=4, mean ± SEM, one-way ANOVA Bonferroni’s multiple comparisons test, \**P*<0.05, \*\**P*<0.01).

As lipid flippase dysfunction is βCTF level-dependent, we investigated the involvement of T-RAP in APP metabolism. After a 48 h T-RAP peptide treatment, the sAPPβ and βCTF levels significantly decreased without altering the full-length APP and TMEM30A protein levels (Fig. 6C and D). Next, we analyzed the T-RAP effect on endosomal morphology in COS-7 cells. Importantly, T-RAP significantly decreased the mean of the Rab5A-positive puncta area in COS-BACE1 cells without altering the total Rab5A protein level (Fig. 6E, F, and Supplementary Fig. 9C). These lines of evidence explain that T-RAP peptide improves endosomal anomalies by rescuing the lipid flippase activity.

## Discussion

βCTF-mediated endosomal anomalies in forming enlarged endosomes are considered early AD pathogenic events^10^. In this study, we showed that lipid flippase activity in endosomes decreased by elevated βCTF level and it contributes to endosomal anomalies. Moreover, in the AD model mice brain, age-dependent lipid flippase dysfunction occurs before Aβ deposition, supporting their strong link with the early pathogenic event. Importantly, we found that a novel βCTF/Aβ-interacting peptide “T-RAP” could recover the lipid flippase activity and endosomal anomalies. Therefore, our findings suggest that lipid flippase impairment is a driving mechanism of βCTF-mediated endosomal anomalies and present a novel therapeutic strategy for AD treatment.

Endosomal anomalies are the signature for vesicular traffic impairment and could be found in the early phase of AD and Down syndrome (DS) prior to Aβ deposition^7^. As DS displays trisomy on chromosome 21, upregulating APP expression, many studies have focused on APP metabolites toxicity. Among these metabolites, the BACE1 product βCTF is considered the driver for endosomal anomalies. Supportively, βCTF accumulation is observed in AD brains^10^, concomitant with upregulated BACE1 expression and activity in AD brains^12^. Several studies using human APP/Presenilin-1 familial AD mutant knock-in iPSC-derived neurons^9^ or 3xTg-AD model mice^8^ have shown that endosome enlargement depends on elevated βCTF but not Aβ. Moreover, analysis of DS fibroblasts or Ts65Dn model mice has demonstrated that accumulated βCTF induces endosome enlargement^29^. However, the underlying mechanisms are not fully explored.

To model the early pathogenic event of AD, we established a BACE1 stably overexpressing neuroblast cell line (SH-BACE1) and observed an increase in BACE1 products, β1CTF (C99) and β11CTF (C89) (Fig. 1A). Interestingly, we found that a subcomponent of lipid flippase, TMEM30A, is the interacting partner for these βCTF (Fig. 1B and Supplementary Fig. 1B). Concomitantly, our organelle fractionation analysis showed a broadened distribution of the endosomal marker protein Rab5A to heavier fractions (Fr 14∼16) in SH-BACE1 cells compared with normal endosome fractions (Fr 7∼12) in control cells. Moreover, TMEM30A, βCTF, and BACE1 were co-distributed in these abnormal heavier fractions (Fig. 1C and Supplementary Fig. 2A). We hypothesize the vicious cycle, in which increased βCTF triggers the interaction with TMEM30A to promote endosomal anomalies, and then the abnormal distribution of BACE1 further promotes βCTF production. In connection with these results, immunofluorescence analysis using COS-7 cells showed the formation of enlarged endosomes in COS-BACE1 cells (Fig. 1D and E). Supportively, our previous study showed that βCTF co-localized with TMEM30A in enlarged endosomes in COS-7 cells^13^. Another study proposed that intracellular Aβ, such as Aβ oligomers, cause vesicular traffic impariment^28^. However, we failed to detect the interaction between TMEM30A and Aβ monomers or oligomers in A7 mice (Supplementary Fig. 7). Although we cannot fully rule out the possibility of the Aβ oligomer involvement in a more advanced stage of AD, our data indicate that the major partner of TMEM30A, which mediated endosome enlargement, is βCTF.

There are two possible mechanisms underlying the TMEM30A/βCTF complex-mediated endosome enlargement. The first is lipid flippase dysfunction and the second is Rab5 overactivation.

Most lipid flippases consist of TMEM30A and active subcomponents, P4-ATPases^30,31^, and regulate phospholipid asymmetry in the lipid bilayer^14^. TMEM30A regulates proper cellular localization and activity of partners, P4-ATPases^24,32^. Intriguingly, we showed the complex formation of TMEM30A and ATP8A1, a brain-enriched endosomal P4-ATPase^31^, decreased in SH-BACE1 cells, and the inhibition of the BACE1 activity recovered the physiological complex formation (Fig. 2A and B). Our data suggest the hypothesis that upregulated BACE1 activity accumulates βCTF, which interacts with TMEM30A to interrupt the complex formation between TMEM30A and ATP8A1. Indeed, the age-dependent disruption of this lipid flippase formation was followed by TMEM30A/βCTF complex formation in AD model mice (Fig. 4A-C). We consider that the TMEM30A/βCTF complex further accumulates βCTF in a vicious cycle, disrupting lipid flippase formation. Supporting this idea, TMEM30A overexpression in COS-7 cells accumulates βCTF, concomitantly with the complex formation of TMEM30A and βCTF^13^.

Endosomal lipid flippases transport phospholipids, like PS, to the cytosolic leaflet^14^. PS on the cytosolic side in endosomes recruits PS-binding proteins, such as Evectin-2, to promote membrane fission and trigger vesicle transport^15,16^. In this study, we focused on Evectin-2 distribution in the endosomes to indirectly monitor lipid flippase activity. We clarified that Evectin-2 distribution decreased depending on the increased βCTF level in SH-BACE1 cells using biochemical analysis (Fig. 2) and the split-luciferase system (Fig. 3). These findings suggest that accumulated βCTF decreases the lipid flippase activity in endosomes to reduce PS level on the cytosolic side. Moreover, the AD model mice analysis indicated decreased Evectin-2 distribution in the membrane fractions (Fig. 4A and D). Indeed, the ATP8A1 knockdown induces endosome-mediated membrane traffic defects^22^. Therefore, we propose the hypothesis that the TMEM30A/βCTF complex impairs lipid flippase formation and its activity to develop endosomal anomalies. It is important to note that our split-luciferase system can be applied to various organelle markers to estimate the localization of proteins and may be developed into a semi-quantitative method for measuring vesicular trafficking. Further validation is needed as a future study.

The second possible mechanism is Rab5 activation. Previous reports, using Rab5A constitutively active mutant (Q79L)^33^ and overexpressing mice^34^, showed Rab5 positive enlarged endosomes. Additionally, elevated βCTF can form a complex with APPL1, a Rab5 effector protein, which mediates Rab5 activation and endosome enlargement in DS fibroblasts and AD brains^10^. Consistent with these reports, we observed the upregulation of BACE1 mediated the abnormal co-distribution of Rab5A and βCTF to alter Rab5A-positive endosomal morphology (Fig. 1C-E). Therefore, accelerated βCTF accumulation by its complex formation with TMEM30A might activate Rab5 to induce endosome enlargement. It would deserve further investigation that lipid flippase dysfunction contributes to Rab5 activation-dependent or independent endosomal anomalies.

Questions still remain concerning how lipid flippase-mediated endosomal anomalies could contribute to AD pathogenesis. Previously, APP-dependent endosomal anomalies and the traffic impairment of nerve growth factor (NGF), which resulted in neuronal atrophy, were observed in rodent neuron models^29^. Another study demonstrated that TMEM30A deficiency influenced sAPPβ and βCTF levels, as well as Aβ/p3 production via the increased β/α-secretase processing of APP^35^. Therefore, lipid flippase dysfunction possibly mediates APP metabolic changes or vesicle transport deficit such as NGF to contribute to AD pathogenesis.

Since vesicular traffic impairment has been implicated as the contributor of AD pathology, βCTF-mediated endosomal anomalies might be promising therapeutic targets for treating AD. The BACE1 inhibitor might be a candidate. According to this notion, β-secretase inhibitor IV recovered lipid flippase function (Fig. 2E, F, and Fig. 3E). However, the study of BACE1 knockout mice showed unexpected neuronal phenotypes, such as schizophrenia endophenotypes or spine density reduction, originating from abrogated β-secretase processing of different substrates such as neuregulin 1 (NRG1)^36,37^. Further therapeutic candidates are pharmacological chaperons to stabilize retromers and limit APP processing in the endosomes by enhancing vesicle transport^38^. There is also a concern that the compound displays a too wide effect range and side effects^39^. These lines of evidence remind us to develop more specific therapeutic targets reflecting AD pathology.

We observed a progressive decline of lipid flippase function in AD model mice (Fig. 4A and D). Moreover, a genome-wide association study identified AD risk variants in one of the P4-ATPases, *ATP8B4*^40^. Therefore, lipid flippase dysfunction could be associated with AD pathogenesis and the optimal target based on AD pathology. Intriguingly, we identified a TMEM30A-derived peptide, “T-RAP,” specifically interacting βCTF and Aβ. T-RAP trended to rescue the lipid flippase activity and improved endosome enlargement in BACE1 stably overexpressing cells (Fig. 6B, E, F, and Supplementary Fig. 9B). We hypothesize that T-RAP enfolds βCTF to inhibit the interaction with TMEM30A, then recovers the lipid flippase physiological formation and activity. This functional lipid flippase recovery could prevent βCTF-mediated endosomal anomalies, enhancing vesicle transport (Fig. 7). T-RAP also decreased sAPPβ and βCTF levels (Fig. 6C and D), which means the peptide corrects vesicle transport to normalize APP processing. Another possibility is that T-RAP directly binds to the proximal site of BACE1 cleavage of APP. Moreover, T-RAP is hydrophilic and easy to handle biochemically. Although further T-RAP specificity analyses would be required, we propose that T-RAP and related molecules might be the optimal candidates for AD therapeutics.

**Figure 7.**
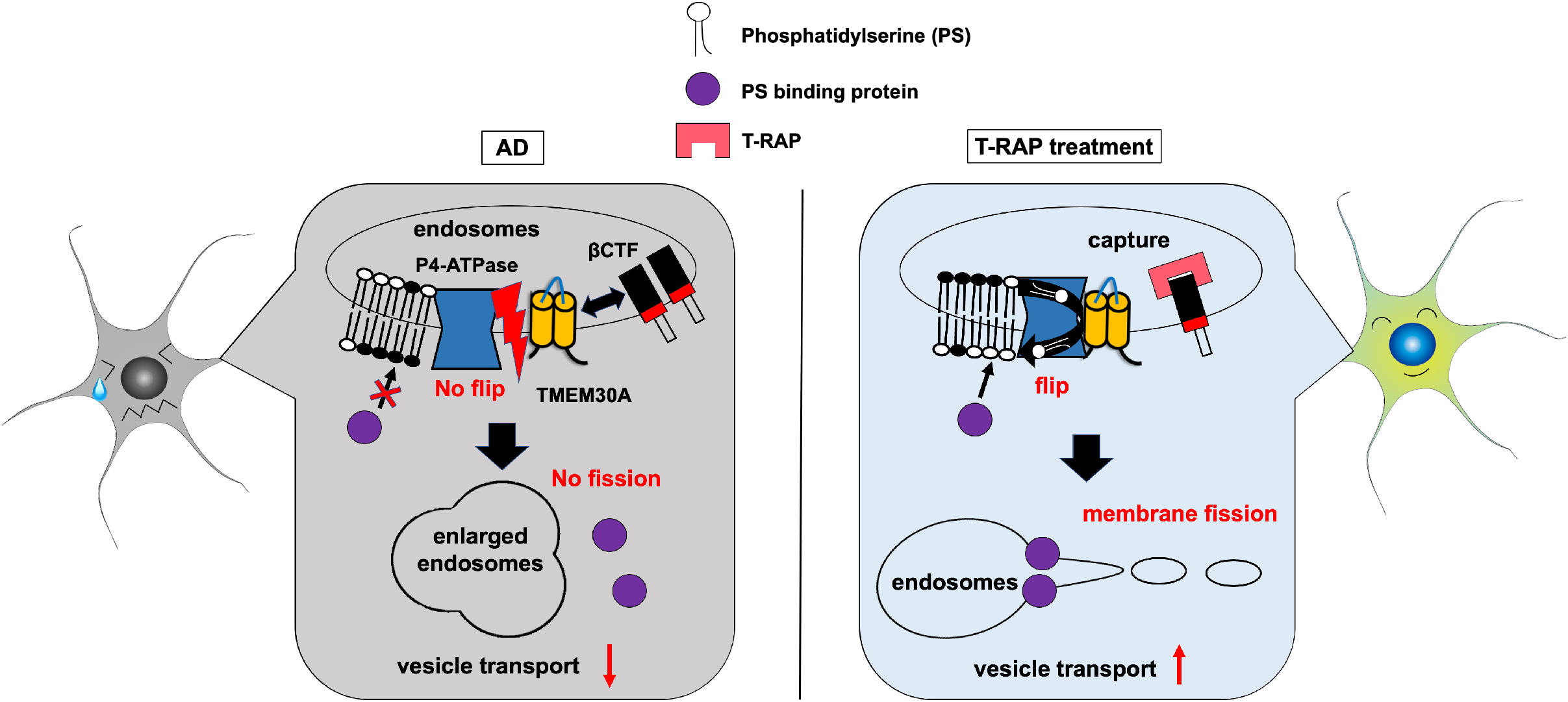
Schematic view of the predicted mechanism of vesicular traffic impairment in AD, and therapeutic effect of T-RAP. In AD, the complex formation between TMEM30A and accumulated βCTF can interrupt lipid flippase physiological formation and its activity in endosomes, decreasing cytosolic PS levels required for membrane fission. Thus, the defect of membrane fission induces endosome enlargement and inhibits vesicle transport. On the other hand, the treatment of βCTF interacting peptide, T-RAP, can prevent the TMEM30A/βCTF interaction, resulting in improved lipid flippase function and endosome enlargement. Therefore, T-RAP treatment can promote vesicle transport.

Three main problems could be distinguished in AD treatment from the perspective of vesicular traffic impairment. First, the details of the mechanisms underlying the endosomal anomalies were unclear. Second, the effective measurement of vesicular trafficking has not been established. Third, endosomal anomaly-related drug treatment has not yet been developed. In this study, we propose lipid flippase dysfunction as a mechanistic contributor for βCTF-mediated endosomal anomalies. Next, our split-luciferase system for measuring endosomal lipid flippase activity could be used for the development of quantitative vesicle transport measurements. Finally, we identified a novel therapeutic candidate for βCTF-mediated endosomal anomalies. Therefore, we present new insights into AD treatment for targeting the early pathogenesis, endosomal anomalies.

## Methods

### Usage of mouse brains

APP transgenic mice (A7 line) have been previously generated^27^ and genotyped using specific primers^41^. All mice were kept under specific pathogen-free conditions and fed a regular diet (Oriental Yeast). The animal care and use procedures were approved by the Institutional Animal Care and Use Committees of the University of Tokyo (18-P-108).

### Statistical analysis

All experimental data are expressed as the mean ± SEM. The experiments were analyzed with two-tailed Student’s t-test or one-way ANOVA with Bonferroni’s multiple comparisons test using the GraphPad Prism8 software. The statistical significance is indicated as follows: \**P* < 0.05, \*\**P* < 0.01, \*\*\**P* < 0.001, \*\*\*\**P* < 0.0001.

## Supplementary information

Detailed information of the materials and methods, original western blot data of Fig. 1-5, as well as the supplementary figures are available in Supplementary information file.

## Acknowledgments

We are grateful to Dr. T. Tomita (University of Tokyo), Dr. N. Nukina (Doshisha University) and Legend research grant in 2019 for providing research materials. This work was supported by the Grant-in-Aid for Scientific Research (C) from the Japan Society for the Promotion of Science (17K08272 and 20K07014), Research Grant from the Ryobi Teien Memorial Foundation, Life Science Foundation of Japan (NT), Grant-in-Aid for JSPS Fellows (NK).

## Author contributions

N.K., and N.T., designed the research; N.K., R.I., M.K., A.I., and N.T. performed the experiments and analyzed data; T.H. and T.S. prepared animal samples and provided antibodies, respectively; N.K., and N.T., wrote the paper; Y.K., T.H., T.I., T.S., and T.U. provided advice and helped write the article. All authors read and approved the final manuscript.

## Conflict of interest

The authors declare that they have no conflict of interest.

## Notes

### Competing Interest Statement

The authors have declared no competing interest.

### Summary of Updates

Correction of author address. Correction of acknowledgement

