## Supplementary Information for "Lipid flippase dysfunction as a novel therapeutic target for endosomal anomalies in Alzheimer’s disease"

#### Materials and Methods

##### Plasmids, antibodies, and reagents

The antibodies, reagents, primers, and synthesized DNA fragments used in this study are listed in supplementary table 1.

*pEB-hyg-BACE1 and pEB-hyg-TO-SC100, -SC89 construction*; Mouse BACE1 sequence in pMx-puro[1] (kind gift from Dr. Gopal Thinakaran, USF Neuroscience Institute), was amplified by PrimeSTAR Max DNA Polymerase (Takara-Bio). PCR products were digested by SalI and BamHI and ligated into the equally digested episomal vector[2], pEB-hyg (Fujifilm-Wako). SC100 sequence was amplified from pcDNA3.1/hygromycin-SC100[3]. The PCR products were introduced into pEB-hyg digested with XhoI and NotI using NEBuilder Hifi DNA assembly mix (New England Biolab) according to the manufacture's instructions. To add Tetracycline inducibility, we digested pEB-hyg-SC100 and pcDNA4 TO (Thermo Fisher) with SnaBI and XhoI and then replaced the CMV promoter with CMV-TO. pEB-hyg-TO-SC89 was produced by PCR mutagenesis using pEB-hyg-SC100 as a template.

*pEB-Puro-HA-LgBiT-mRab5;* To generate N-terminally HA-tagged Large BiT sequence (HA-LgBiT), we performed PCR reaction twice. At 1$^{st}$ round, the LgBiT sequence was amplified, using CMV HaloTag-LgBiT (Promega) as a template. Using 1$^{st}$ PCR product as a template, 2$^{nd}$ PCR was performed to enable the Gibson assembly reaction. Murine Rab5A (mRab5A) was amplified from DEST40-CFP-mRab5A. PCR products, HA-LgBiT and mRab5A, were assembled into pEB-puro digested with XhoI and BamHI using NEBuilder Hifi DNA assembly mix.

*p3xFLAG CMV10-SmBiT-Evectin-2 / -K20E mutant*; To generate Evectin-2 N-terminally fused with Small BiT (SmBiT) fragment, we amplified the SmBiT-Evectin-2 sequence, using pEGFP-N1-Evectin-2 embedding human Evectin-2 CDS as a template. The SmBiT sequence was obtained from Promega. Using the PCR product as a template, we amplified the insert sequence to enable the Gibson assembly reaction. The PCR product inserted into the p3xFLAG-CMV10 digested with EcoRI and NotI using NEBuilder Hifi DNA assembly mix. K20E mutant was produced by long-PCR mutagenesis using p3xFLAG CMV10-SmBiT-Evectin-2 as a template.

*pENTR-U6-shRNA vectors*; The U6 promoter sequence was synthesized by Thermo Fisher Scientific and introduced into pENTR/D-TOPO (Thermo Fisher Scientific). Indicated shRNA template pairs were annealed and ligated into pENTR-U6 digested by AgeI and EcoRI.

*pDEST-15-constructs*; The codon sequence of the extracellular region of TMEM30A (67-323 AA: TmEx) was redesigned to minimize the usage of the rare codons in E. Coli and synthesized (Tm-EXN; Genscript). To stepwise shortening of the TmEx domain, PCR amplification was performed using primers listed in Sup using Tm-Ex sequence as a template. Each PCR product was cloned into the pENTR/D-TOPO vector. The target sequence was subcloned into pDEST-15 (Thermo Fisher Scientific) to make GST fusion protein.

Resrictiction enzymes used in this study were purchased from New England Biolabs.

All constructed plasmids were verified the inserted sequence with Sanger sequencing (GENEWIZ).

##### Cell culture

The human neuroblastoma SH-SY5Y cell or the African green monkey kidney fibroblast COS-7 cell was transfected with indicated plasmids to establish stably overexpressing cell lines. Subsequently, these cells were selected by hygromycin (50 mg/ml, Fujifilm-Wako). Human embryonic kidney cell line HEK293 or HEK293T and COS-7 cells were cultured at 37°C with 5% $CO_2$ in Dulbecco's modified Eagle's medium (DMEM, Fujifilm-Wako), and SH-SY5Y cells were maintained in DMEM/Ham's F-12 (Fujifilm-Wako). Media were supplemented with 10% fetal bovine serum, 100 units/ml penicillin, and 100 μg/ml streptomycin.

**Transfection and reagents treatments**

Cells at 50~70% confluence were transfected with indicated plasmids by Lipofectamine 3000 Transfection Reagent (Thermo Fisher Scientific) following the manufacture's protocol. For the interaction assay for TMEM30A and β1 or β11CTF, HEK293T cells were co-transfected with CFP-TMEM30A and SC100 (β1) or SC89 (β11), then applied to co-immunoprecipitation analysis after 48 h from the transfection. For the knockdown of TMEM30A, cells were reversely transfected with short hairpin RNA for each target inserted in the pENTR-U6 vector and incubated for 72 h. For the knockdown of APP, siRNA (SASI_Hs01_00185800, Sigma) was reversely transfected by Lipofectamine RNAiMAX Transfection Reagent (Thermo Fisher Scientific) following the manufacture's protocol and maintained for 72 h at 37°C. Cells were treated with 10 μM of β-secretase inhibitor IV or T-RAP and 50 μM of TAT-T-RAP for 48 h in all experiments.

**Alamar Blue assay**

To monitor the T-RAP toxicity, we performed Alamar Blue analysis (Bio-rad) according to the manufacture's instruction. After 48 h treatment of T-RAP (10 uM) or ethanol in SH-SY5Y cells, the medium was replaced with Almar Blue containing medium. After 1 h incubation, medium fluorescence was measured at 530 nm excitation and 590 nm emission wavelengths.

**Usage of mouse brains**

The transgenic mice that overexpress human APP695 harboring K670N, M671L, and T714I FAD mutations in neurons under the control of Thy1.2 promoter (A7 line) were generated previously[4] and genotyped using specific primers[5]. All mice were kept under SPF conditions and fed a regular diet (Oriental Yeast). The animal care and use procedures were approved by the Institutional Animal Care and Use Committees of the University of Tokyo (18-P-108). Whole brains were removed from the skull, and brain hemispheres of female A7 mice at 3- or 6-month-old, and age-matched controls were prepared. Tissue samples were stored at -80°C until biochemical analysis.

**Preparation of membrane fractions from cells and mouse brains**

Cells were washed with cold phosphate-buffered saline (PBS), then homogenized in buffer A (10 mM HEPES pH7.4, 150 mM NaCl, 10% glycerol) by Dounce homogenization and 21G syringe at 20 strokes. After centrifugation at 1,500 g, 10 min, 4°C, the supernatant was collected. The ppt was resuspended with buffer A and repeated the solubilization step. Then, resultant supernatant was combined previous one as post-nuclear supernatant (PNS). Crude membrane fraction (MF) was obtained from PNS by ultracentrifugation at 100,000 g, 1 h, 4°C in an SW41Ti rotor (Beckman Coulter). To monitor Evectin-2 localization in membranes, MF was dissolved in RIPA buffer (50 mM Tris-HCl pH 8.0, 150 mM sodium chloride, 0.5% sodium deoxycholate, 0.1% sodium dodecyl sulfate, 1% NP-40). To analyze the lipid flippase complex formation, MF was dissolved completely in buffer A containing 1% CHAPS by 21G syringe at 20 strokes and rotated for 1 h at 4°C. After excluding the insoluble fractions by centrifugation at 15,000 rpm, 10 min, 4°C, the supernatants were performed in co-immunoprecipitation. For mouse brain samples, a cerebral hemisphere was homogenized in buffer B (0.32 M sucrose, 10 mM HEPES pH 7.4, 1 mM EDTA) by Dounce homogenization at 15 strokes. PNS was collected after centrifugation at 1,500 g, 10 min, 4°C at 2 times. The crude membrane isolation step of mouse brain samples was the same as described above, then MF was dissolved in buffer C (50 mM HEPES pH 7.4, 150 mM NaCl, 1% CHAPSO). The supernatants after centrifugation were used for co-immunoprecipitation.

**Co-immunoprecipitation**

Cells were dissolved in IP buffer (20 mM HEPES pH 7.4, 150 mM NaCl, 0.1 mM EDTA, 1% CHAPS) for 30 min at 4°C, and total lysates were clarified centrifugation at 15,000 rpm, 10 min, 4°C. The supernatants were rotated with the indicated antibody for 2 h at 4°C. Then, equilibrated protein G sepharose 4 fast flow beads (GE Healthcare) were added to them and incubated overnight at 4°C. The membrane fractions from cells or mouse brains were performed in the same protocol as the total cell lysates, except for solubilizing buffer condition.

##### GST pull-down

For GST pull-down assays, cells were lysed in 1% CHAPS lysis buffer (50 mM HEPES pH 7.4, 150 mM NaCl, 1 mM EDTA supplied with protein inhibitor cocktail (Roche)), 4°C for 30 min. Lysates were clarified by centrifugation for 5 min at 12,000 g and incubated with GST-fusion proteins pre-bound to GSH-agarose beads (GE Healthcare) for 2 hr at 4°C. The Laemmli sample buffer was added to each precipitate, which was subjected to immunoblotting analysis.

##### Iodixanol gradient fractionation

Subcellular fractionation was conducted as previously described[6]. Briefly, cells were homogenized in HB (250 mM Sucrose, 20 mM Tris-HCl pH 7.4, 1 mM EGTA, 1 mM EDTA) by Dounce homogenization and passing through 21G syringe. PNS was adjusted to 25% iodixanol by mixing with 50% iodixanol in HB. 50% iodixanol was prepared by 60% iodixanol solution (optiprep, Cosmo Bio). 2 ml of 25% mixture was placed at the bottom of 13.2 ml ultracentrifuge tube (Beckman Coulter) and gently overlaid with 1 ml of 20, 18.5, 16.5, 14.5, 12.5, 10.5, 8.5, 6.5, and 5% iodixanol in HB, respectively. The gradient was ultracentrifuged at 27,000 rpm (124,806 g), 20 h, 4˚C in an SW41Ti rotor. Fractions were subsequently collected from the top, then performed TCA precipitation (Trichloroacetic acid, Fujifilm-Wako). After TCA precipitation, pellets were diluted in RIPA buffer and 2X Laemmli sample buffer, then analyzed by immunoblotting.

##### Immunoblotting

For immunoblotting analysis, aliquots of cell lysates were separated on 10 or 12% Tris-glycine gels as previously described[7]. To segregate each APP-CTF and Aβ, high-resolution electrophoresis was performed using 16% Tris-Tricine gels. The separated proteins on PVDF membranes were incubated with appropriate primary antibodies overnight at 4˚C, and horseradish peroxidase secondary antibodies were applied for 1 h at room temperature. Membranes were detected by ImmunoStar LD or Zeta (Fujifilm-Wako), then visualized by the ChemiDoc MP Imaging System (Bio-Rad). The quantification analysis of target proteins was performed using Image Lab software (Bio-Rad).

##### Immunofluorescence

Cultured cells on glass coverslips were fixed in 4% paraformaldehyde / PBS for 20 min and permeabilized with 0.1% Triton X-100 / PBS for 10 min, then wash with PBS and blocked with 1% BSA / PBS for 1 h. Coverslips were incubated with primary antibodies overnight at 4˚C. After washing with 0.05% tween / PBS, coverslips were incubated with secondary antibodies for 30 min at room temperature, washed with 0.05% tween / PBS, and mounted with Vectashield with 4′,6-diamidino-2-phenylindole (Vector Laboratories, Inc). Antibodies are listed in supplementary table 1. Stained cells were imaged on microscope BZ-X810 (Keyence) using the manufacture's software. For Rab5 positive endosome analysis, to increase the resolution and confirm particle focusing, the z-stack images were captured at 0.1 μm intervals along the z-axis using a 60x objective oil lens, and 25 z-stack images were merged into one full-focus image with BZ-X Analyzer software. The 10 fields of images were randomly selected in each sample. To measure the Rab5A positive puncta size and particle distribution, all captured images were converted to binary data and automatically analyzed under the same condition using BZ-X Analyzer Hybrid Cell Count and Macro Cell Count software (Keyence)[8,9]. The average size per particle shows the mean of Rab5A positive puncta area. The size distribution of Rab5A positive puncta has been presented as the percentage of the total particle counts. In three experiments, about 60~90 cells in each sample were analyzed. For T-RAP treatment, about 100~300 cells per sample were analyzed in four experiments.

**Split-luciferase assay**

Cells were cultured in 24 well plates (VisiPlate-24 TC 24well, PerkinElmer) and transfected proteins fused with luciferase subunits, SmBiT-Evectin-2 and LgBiT-mRab5, at 60~70% confluence. For the effect of WT or K20E mutant on lipid flippase activity, cells were reversely transfected with plasmids and measured 48 h after transfection. For the treatment of BACE1 inhibitor, cells were reversely transfected with plasmids, then treated with β-secretase inhibitor IV after 24 h from the transfection. Luciferase activity was measured 48 h after treatment. For the knockdown of TMEM30A, the luciferase activity was measured 72 h after the reverse co-transfection of shRNA-TMEM30A and luciferase subunits plasmids. For the knockdown of APP, cells were transfected with luciferase subunits 24 h after the knockdown of APP. The luciferase activity was measured 72 h after the siRNA transfection. The luminescent signal was obtained by GloMax® Discover (Promega) using the Nano-Glo Live Cell Assay System (N2011, Promega) following the manufacture's protocol. After measurement, cells were lysed with RIPA buffer and performed the BCA protein assay. RLU (Relative Light Unit) except for background was set as follows. The obtained signals were normalized by protein concentration against the background sample (empty vector) to minimize transfection stress. Next, these values were normalized by the ratio of transfection, the expression level of SmBiT-Evectin-2 quantified by immunoblotting against the control sample. Background RLU means the obtained signal. In statistical analysis, RLU with background subtracted was applied.

$$\text{RLU= obtained signal X} \underbrace{\boxed{\frac{\text{Other samples (luciferase subunits)}}{\text{background (empty vectors)}}}}_{\text{Ratio of protein concentration}} \text{ X } \underbrace{\boxed{\frac{\text{SmBiT-Evectin-2 level in each sample}}{\text{SmBiT-Evectin-2 level in control}}}}_{\text{Ratio of transfection}}$$

**Aβ immunoprecipitation by 4G8 antibody**

Aβ levels in the culture media were analyzed as previously described[10]. Briefly, conditioned media added 0.01% Triton X-100, 4G8 antibody, protein A / G agarose beads (Santa Cruz Biotechnologies), and then rotated overnight at 4˚C. After centrifugation at 4,000 rpm, 3 min, 4˚C, pellets were mixed with 4G8 wash buffer (50 mM NaCl, 10 mM Tris-HCl pH 7.6, 0.01% Triton X-100) 20 min at 4˚C. After centrifugation at 4,000 rpm, pellets mixed with 4G8 wash buffer were carefully layered on 1 ml of 1M sucrose in 4G8 wash buffer, then centrifuged at 10,000 rpm, 1 min, 4˚C. Pellets were added 10 mM Tris-HCl pH 7.6 and gently vortexed. After centrifugation at 4,000 rpm, 3 min, 4˚C, pellets were resuspended in 2X Laemmli sample buffer, and samples were boiled immediately before immunoblotting.

**Preparation of synthetic Aβ oligomer**

Aβ42 oligomers were prepared as previously described[11]. Briefly, human synthetic Aβ42 peptide dissolved in 1,1,1,3,3,3-hexafluoro-2-propanol (HFIP, Sigma) was evaporated then resolved in dimethyl-sulfoxide (DMSO) as 1 mg/ml stock at -20˚C. Before use, Aβ42 in DMSO was sonicated in an ultrasonic bath sonicator for 10 min then diluted in DMEM/F12 phenol red-free (Fujifilm-Wako) at 100 μM and left for one day at 4˚C.

**Statistical analysis**

All experimental data are expressed as the mean ± SEM. The experiments were analyzed with a two-tailed Student's t-test or one-way ANOVA with Bonferroni's multiple comparison test using the GraphPad Prism8 software. The statistical significance is indicated as follows: $^{*}P < 0.05$, $^{**}P < 0.01$, $^{***}P < 0.001$, $^{****}P < 0.0001$.

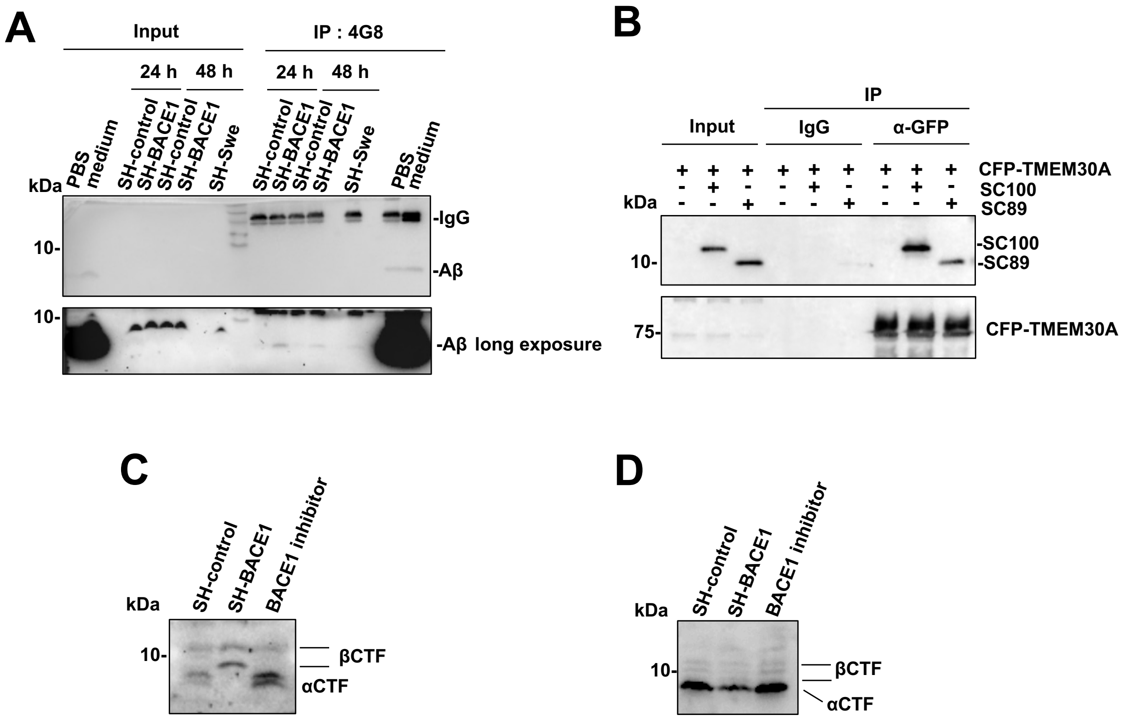


**Supplementary Figure 1. Analyzing for secreted Aβ and the interaction between TMEM30A and βCTF.** (**A**) Immunoprecipitation analysis for secreted Aβ in the culture medium of SH-control and SH-BACE1 cells at 24 h and 48 h after cell seeding using 4G8 antibody. Synthetic Aβ40 in PBS or medium was used as the positive control for immunoprecipitation. Conditioned medium from stably overexpressing APP (Swedish) cells was used as the positive control for increasing Aβ levels. Aβ was detected by 82E1 antibody. (**B**) Co-immunoprecipitation analysis using GFP antibody 48 h after the co-transfection of CFP-TMEM30A, SC100 (β1), and SC89 (β11) in HEK293T cells. SC100, SC89, or CFP-TMEM30A were detected by APP C terminal antibody or TMEM30A antibody, respectively. (**C**) APP-CTF levels in SH-control and SH-BACE1 cell lysate solubilized in RIPA buffer, the same data as Fig.1a for comparison to (d). (**D**) APP-CTF levels in membrane fractions solubilized in 1% CHAPS/buffer A (see supplementary methods).

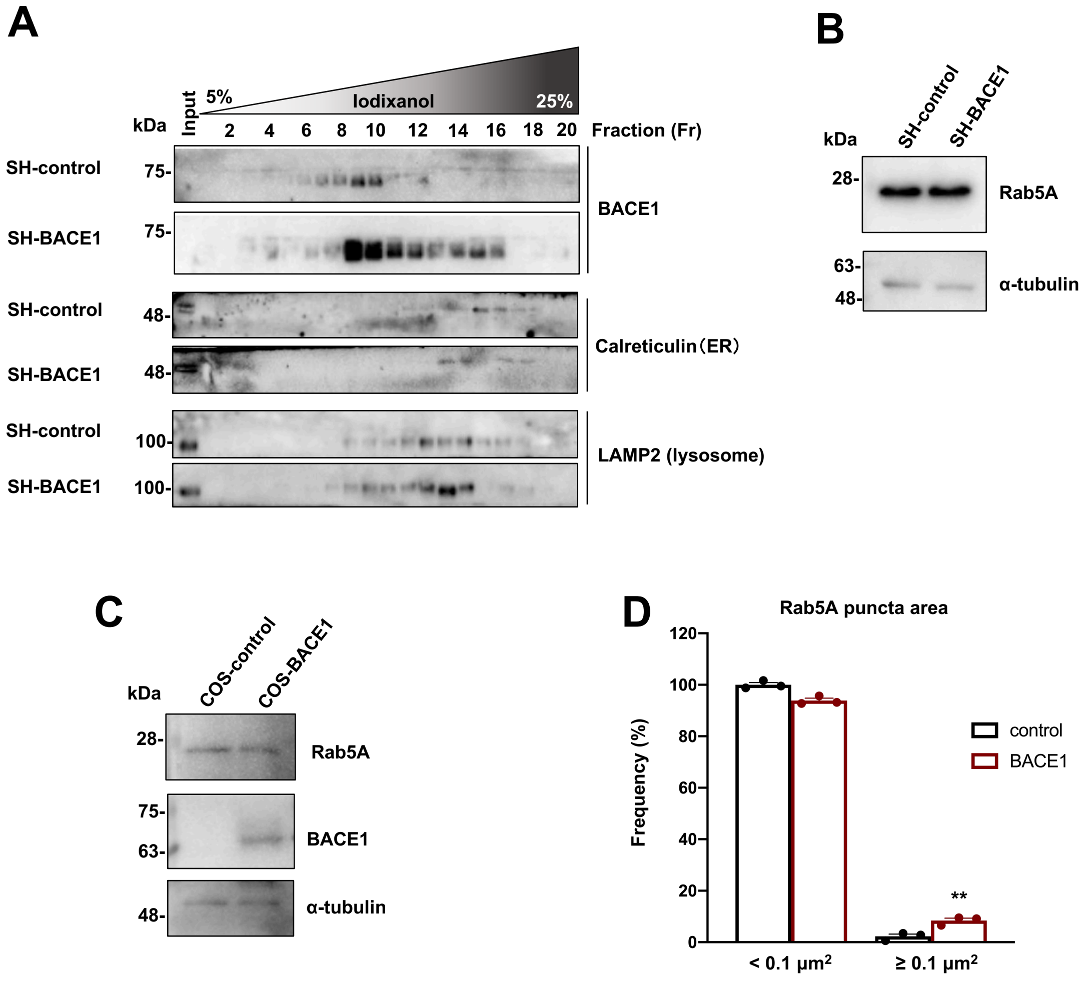


**Supplementary Figure 2. Iodixanol gradient fractionation and immunofluorescence analysis of SH-control and SH-BACE1 cells.** (**A**) The homogenates from SH-control and SH-BACE1 cells were applied to iodixanol gradient fractionation. Calreticulin or LAMP2 were detected as organelle markers for endoplasmic reticulum (ER) or lysosome, respectively. (**B**) Immunoblotting analysis for Rab5A in whole cell lysates of SH-control and SH-BACE1 cells. (**C**) Immunoblotting analysis for Rab5A and BACE1 in whole cell lysates of COS-control and COS-BACE1 cells. (**D**) Quantification of the size distribution (<0.1 μm$^2$ and ≥ 0.1 μm$^2$) of Rab5A positive puncta related to Fig. 1E ($n$=3, mean ± SEM, two-tailed Student's t-test, **$P$<0.01).

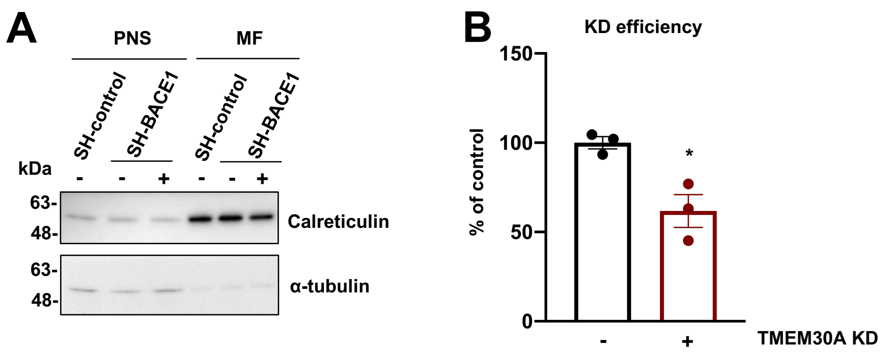


**Supplementary Figure 3. Membrane fractionation and knockdown efficiency of TMEM30A.** (**A**) Calreticulin and α-tubulin were used to confirm the efficacy of membrane fractionation in Fig. 2A. (**B**) Quantification of TMEM30A knockdown in Fig. 2C ($n$=3, mean ± SEM, two-tailed Student's t-test, *$P$<0.05).

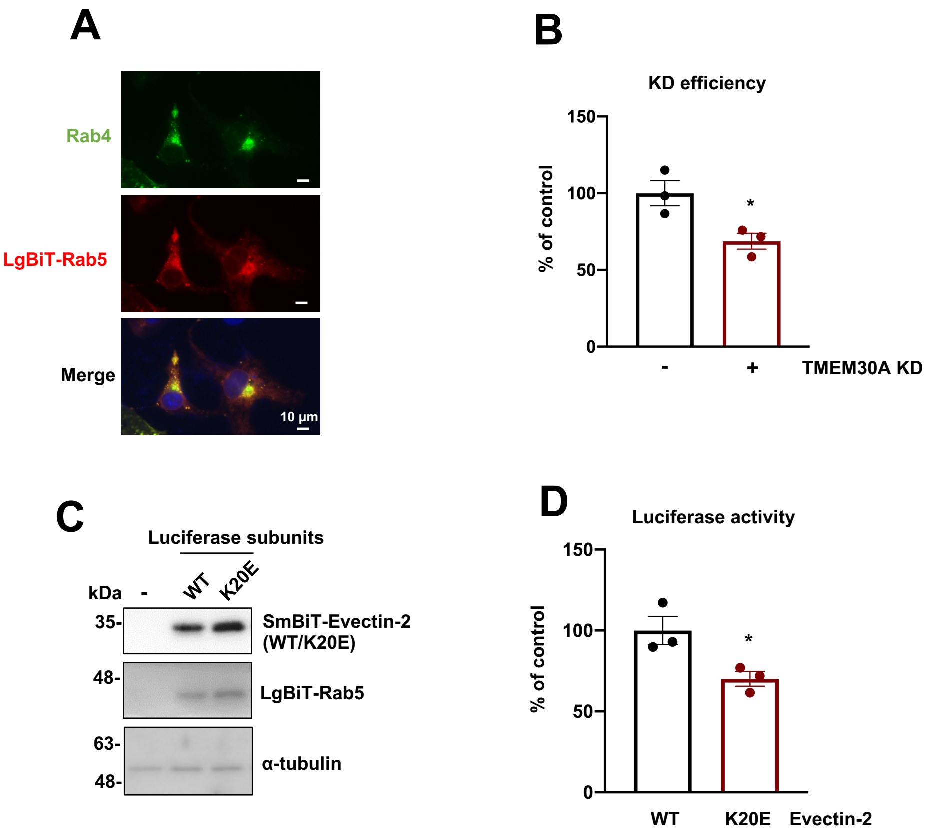


**Supplementary Figure 4. Validation of split-luciferase assay.** (**A**) Confirmation of LgBiT-HA-Rab5 distribution in endosomes. COS-7 cells were transfected with LgBiT-Rab5, after 24 h transfection, and immunostained with endosome marker Rab4 and HA antibodies. Scale bars: 10 μm. Representative images were captured using a 60x objective lens (zoom x1.7). (**B**) Quantification of TMEM30A in Fig. 3B ($n$=3, mean ± SEM, two-tailed Student's t-test, *$P$<0.05). (**C**) Immunoblotting analysis for LgBiT-Rab5, SmBiT-Evectin-2, and -K20E mutant in COS-7 cells. (**D**) Quantification of the luciferase activity in COS-7 cells after 24 h transfection of WT or K20E mutant ($n$=3, mean ± SEM, two-tailed Student's t-test, *$P$<0.05).

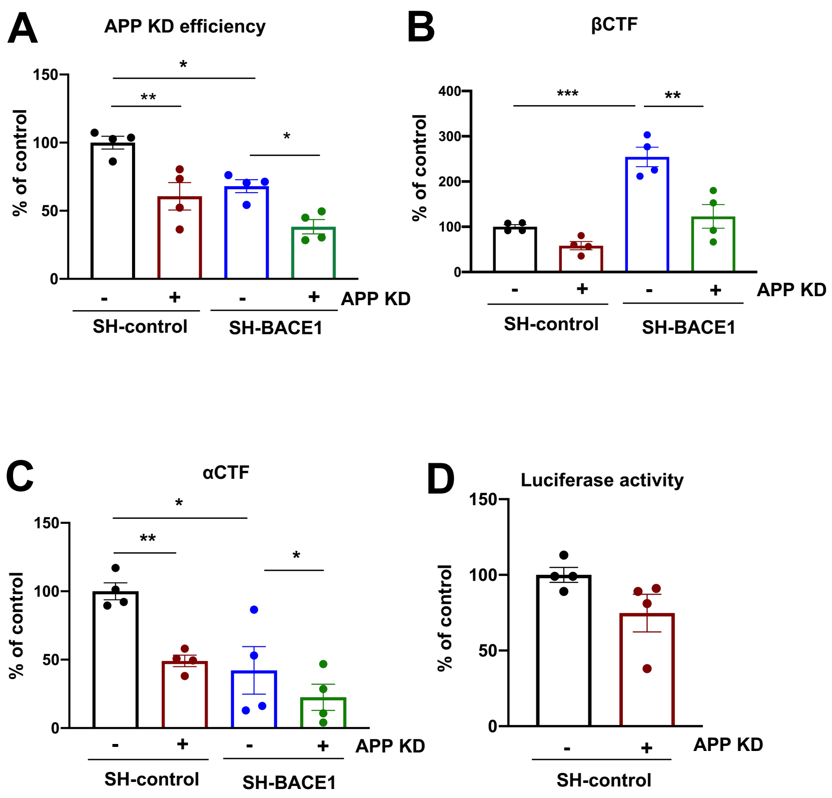


**Supplementary Figure 5. Protein levels of APP metabolites and the luciferase activity of SH-control cells in the knockdown of APP.** (**A-C**) Quantification of APP and APP-CTF in Fig. 3F (*n*=4, mean ± SEM, one-way ANOVA Bonferroni's multiple comparisons test, **P*<0.05, ***P*<0.01, ****P*<0.001). (**D**) Quantification of luciferase activity at 72 h after the knockdown of APP in SH-control cells (*n*=3, mean ± SEM, two-tailed Student's t-test).

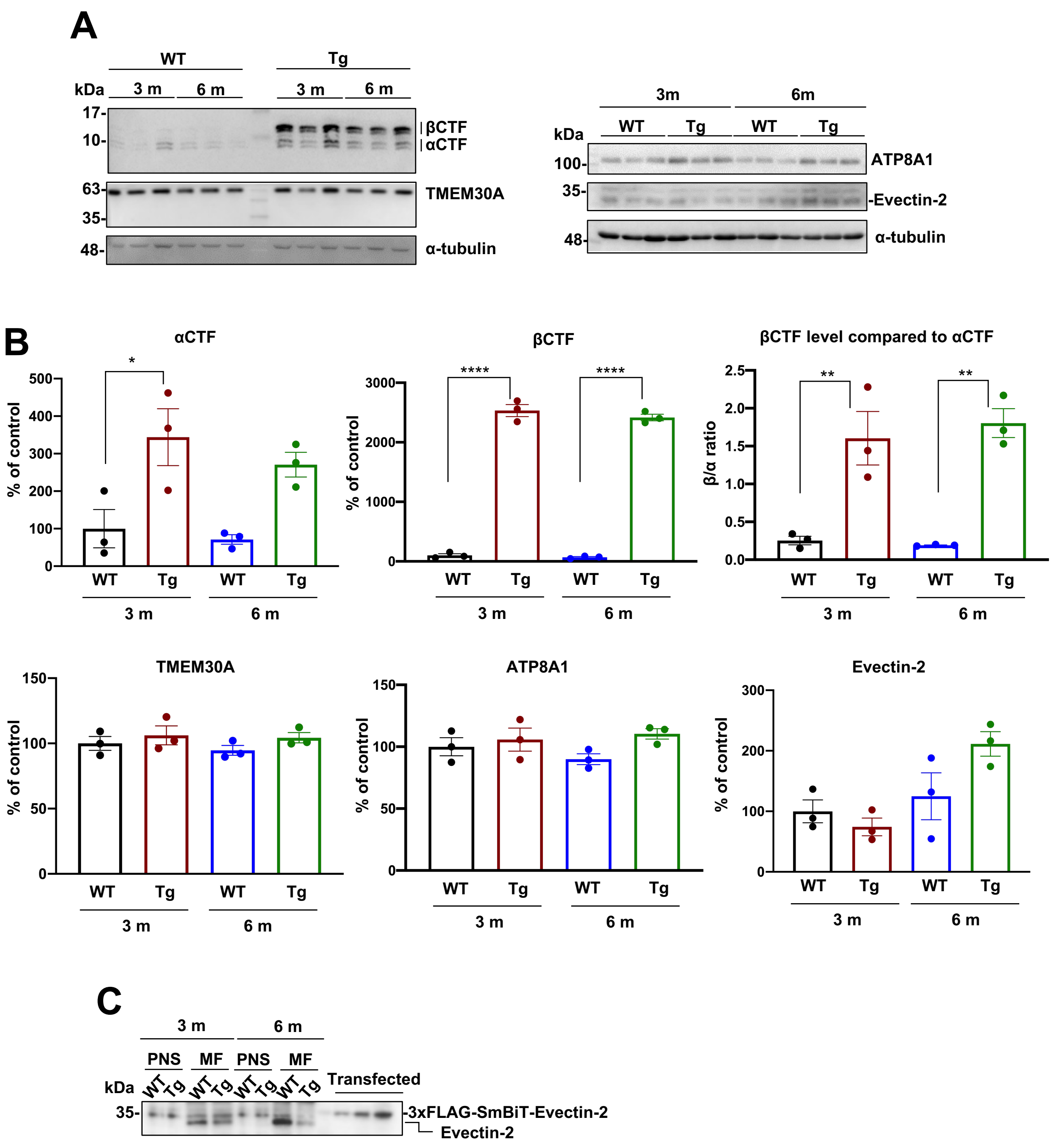


**Supplementary Figure 6. Protein levels in WT and A7 mice at 3 or 6 months.** (**A**) Immunoblotting analysis for APP-CTF, TMEM30A, ATP8A1, and Evectin-2 in WT and A7 mice at 3 or 6 months. (**B**) Quantification of APP-CTF, the ratio of β/α, TMEM30A, ATP8A1, and Evectin-2 in Fig. S6A ($n$=3, mean ± SEM, one-way ANOVA Bonferroni's multiple comparisons test, *$P$<0.05, **$P$<0.01, ****$P$<0.0001). (**C**) Confirmation of endogenous Evectin-2 molecular weight. 3xFLAG SmBiT-Evectin-2 (35 kDa) was used as the positive control.

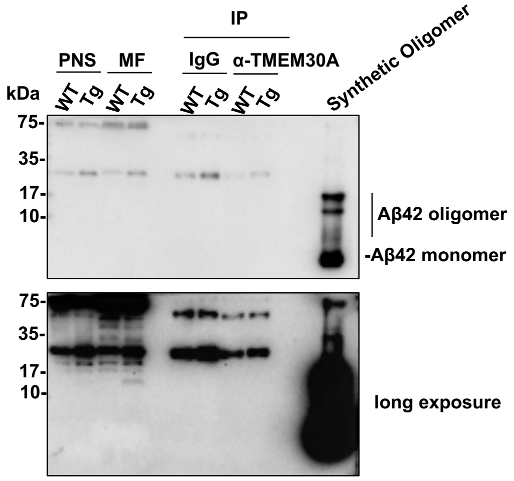


**Supplementary Figure 7. TMEM30A fails to interact with Aβ monomers and oligomers in A7 mice.** The membrane fractions from WT and A7 mice brains at 6 months were performed co-immunoprecipitation analysis using TMEM30A antibody. Aβ monomers and oligomers were detected by 6E10 antibody.

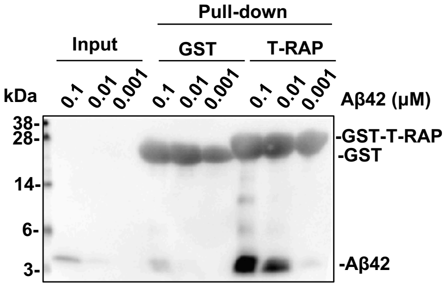


**Supplementary Figure 8. T-RAP interacts with Aβ.** GST-pull down from synthetic Aβ42 at the indicated concentration.

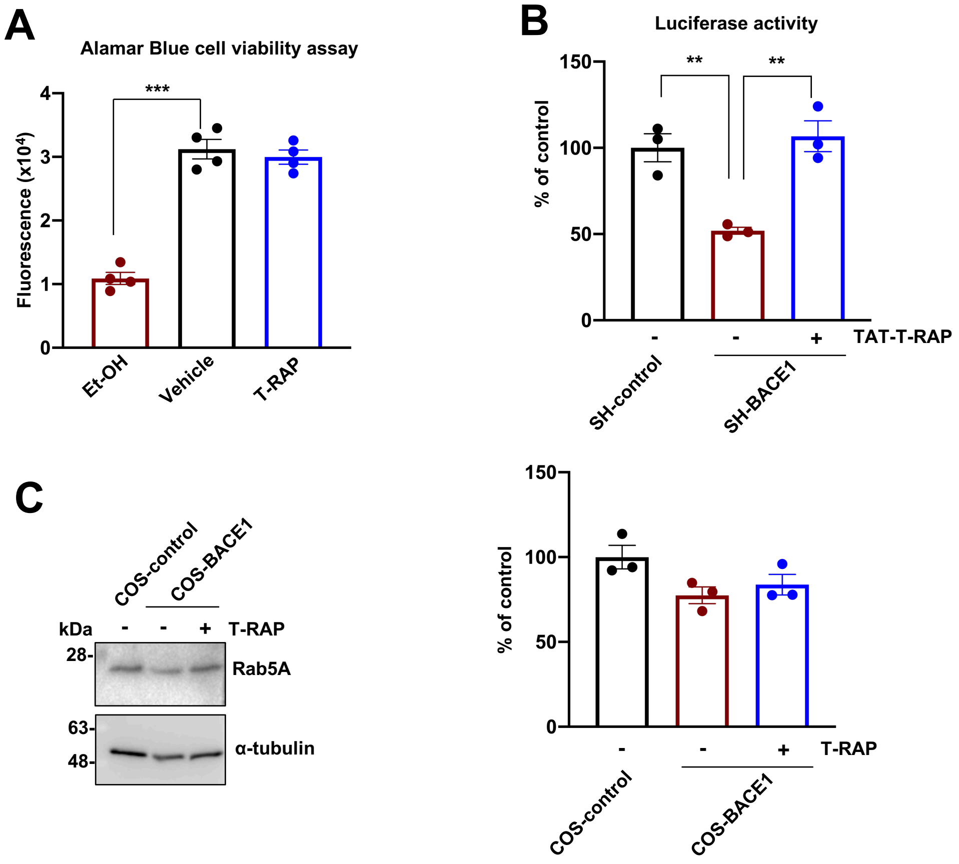


**Supplementary Figure 9. Effects of T-RAP on cell viability, lipid flippase activity, and Rab5 protein levels.** (**A**) Quantification of Alamar Blue cell viability assay in SH-SY5Y cells (*n*=4, mean ± SEM, one-way ANOVA Bonferroni's multiple comparisons test, ***$P < 0.001$). Ethanol was used as a control for cell toxicity. (**B**) Quantification of the luciferase activity after 48 h treatment of TAT-T-RAP (50 μM) (*n*=3, mean ± SEM, one-way ANOVA Bonferroni's multiple comparisons test, **$P<0.01$). (**C**) Immunoblotting analysis for Rab5A in whole cell lysates of COS-control and COS-BACE1 cells after 48 h treatment of T-RAP (10 μM), and quantification data (*n*=3, mean ± SEM, one-way ANOVA Bonferroni's multiple comparisons test).

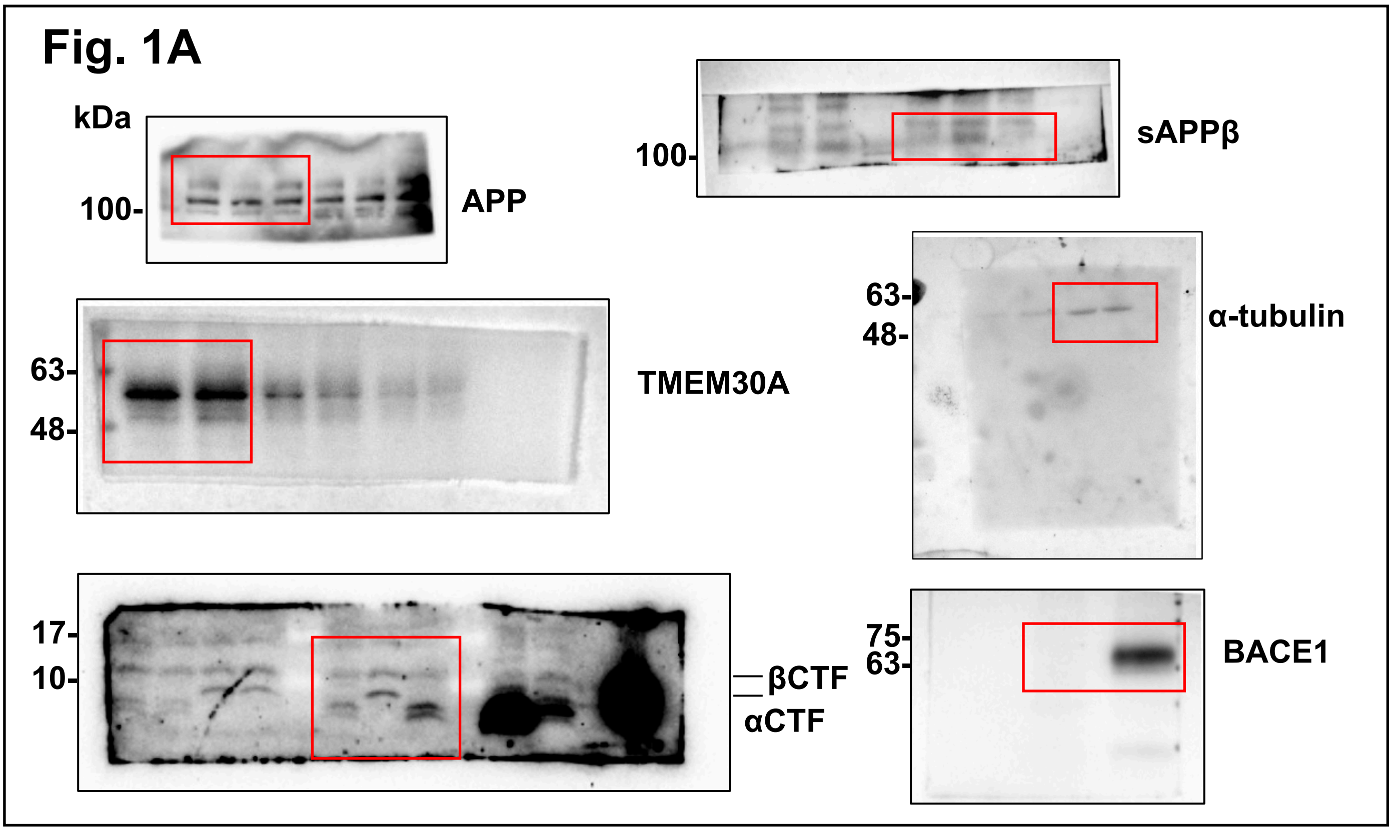


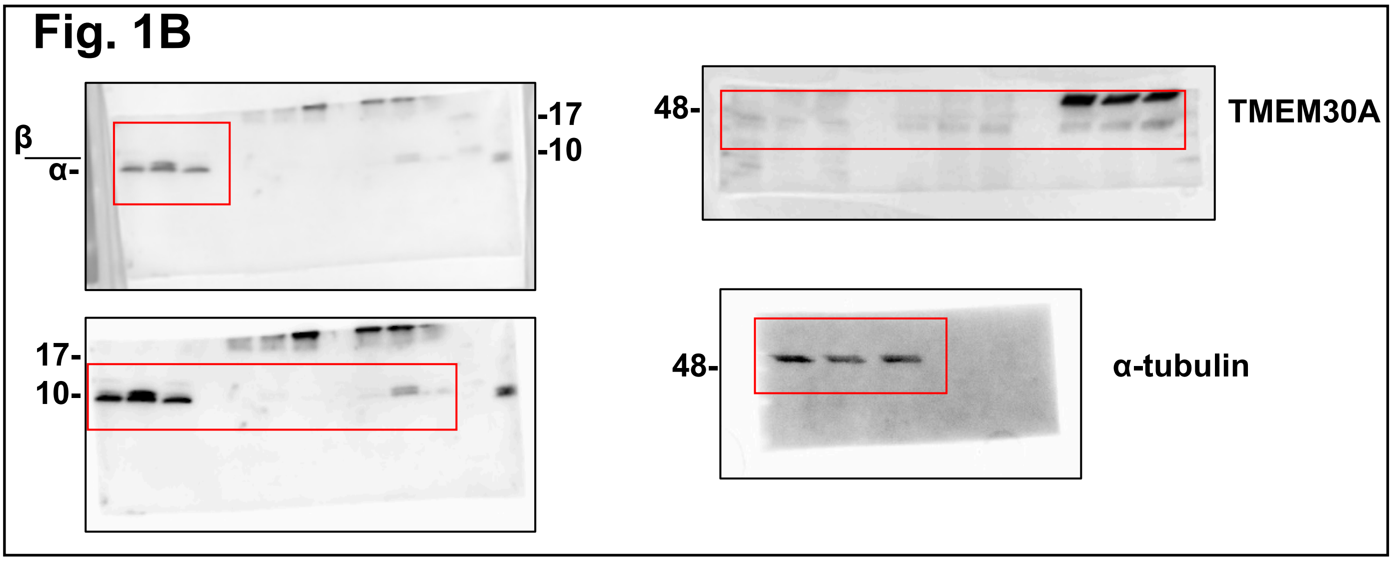


**Supplementary Figure 10. The original western blot data of Fig. 1~6.**

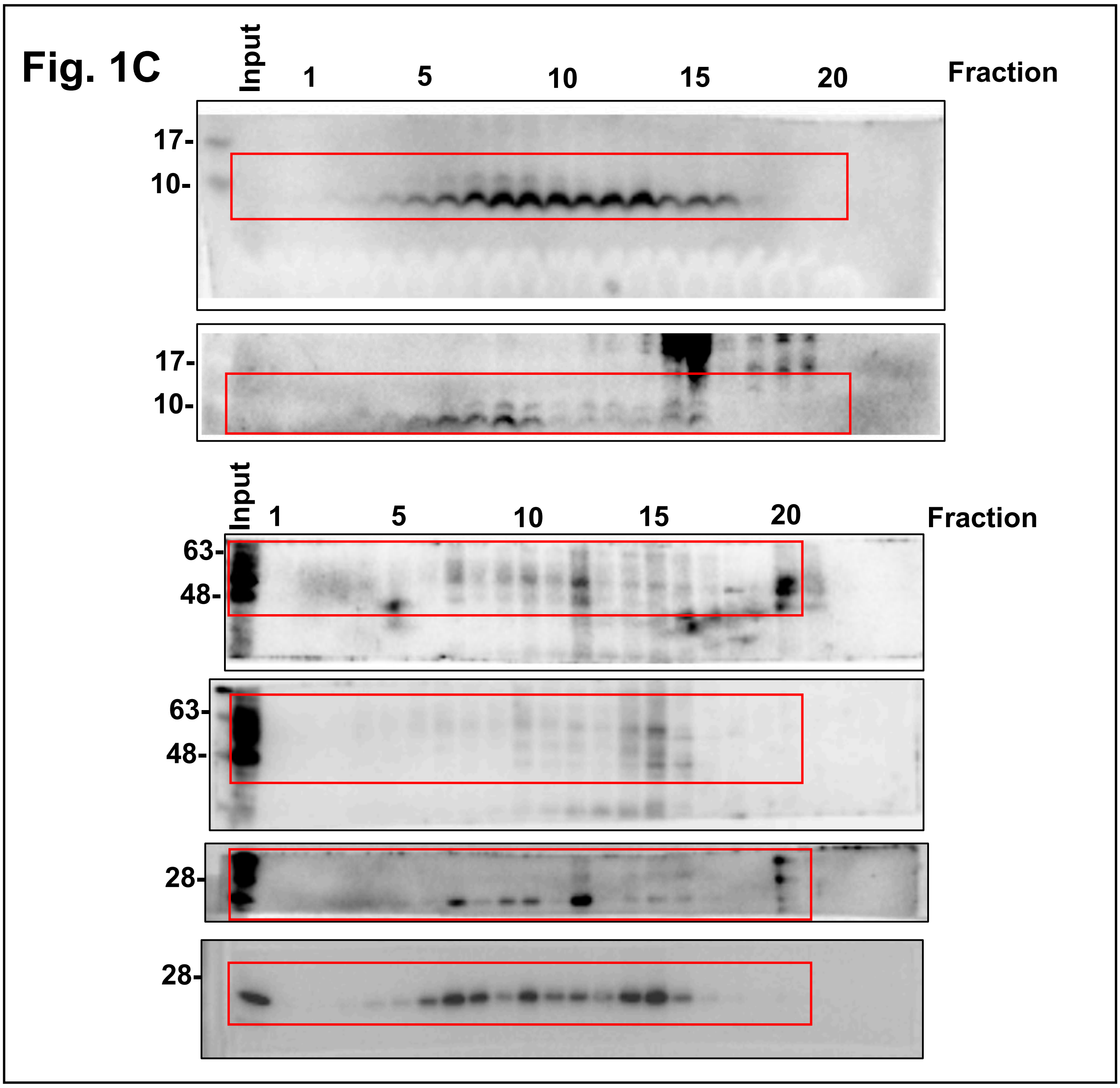


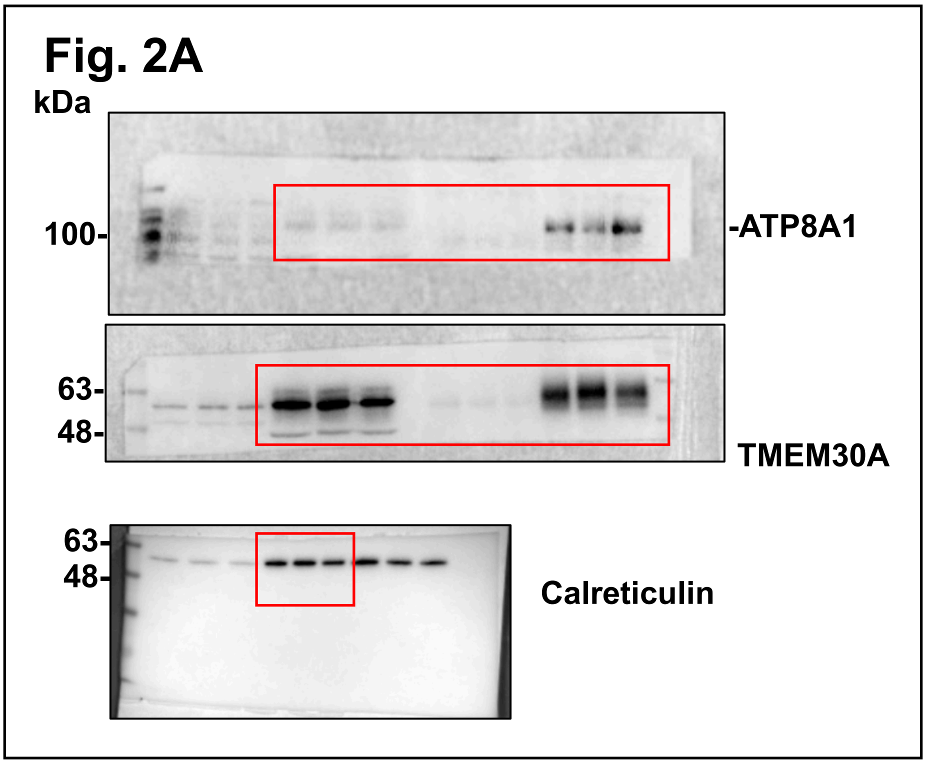


Supplementary Figure 10 (continued). The original western blot data of Fig. 1~6.

#### Fig. 2C

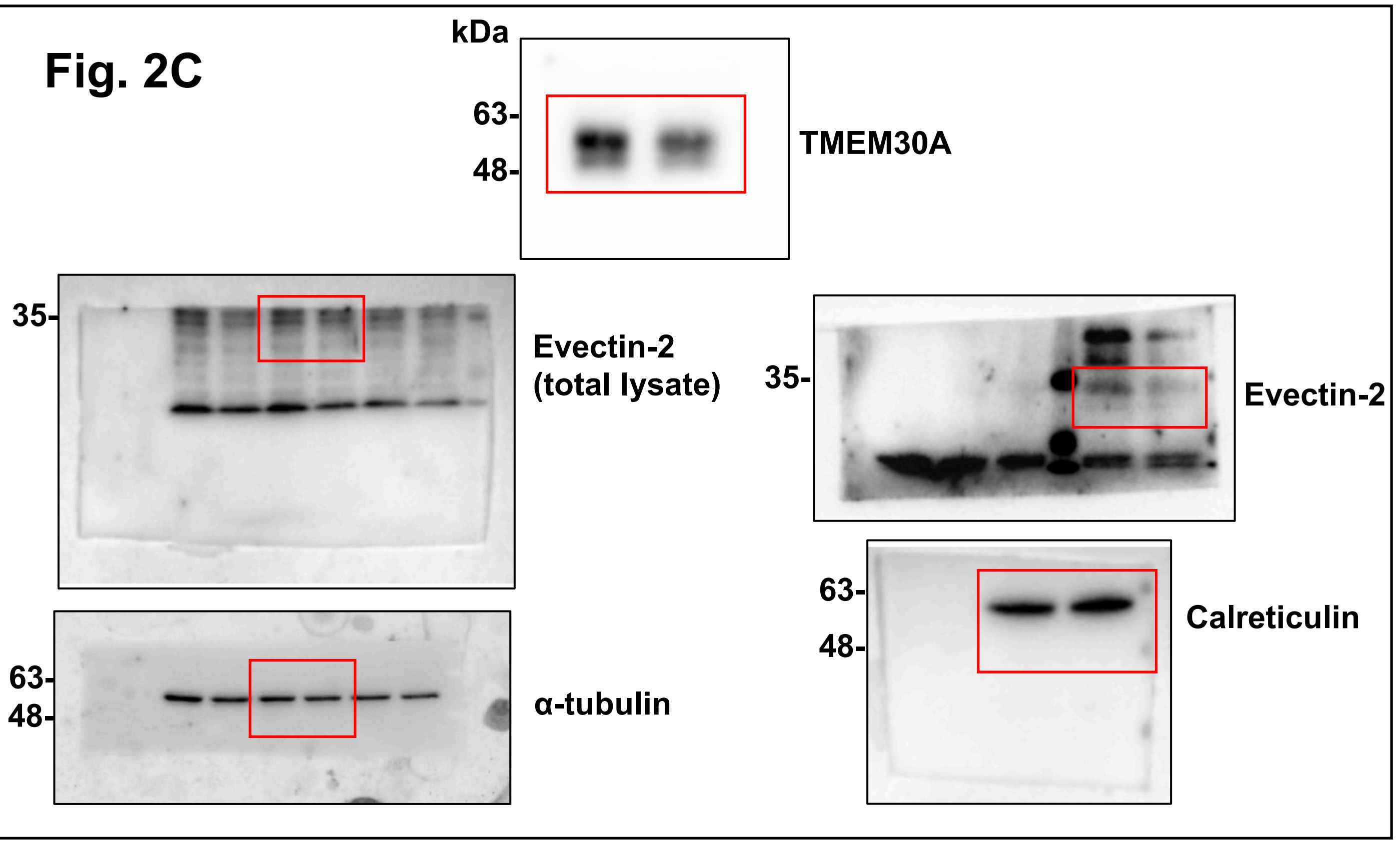


#### Fig. 2E

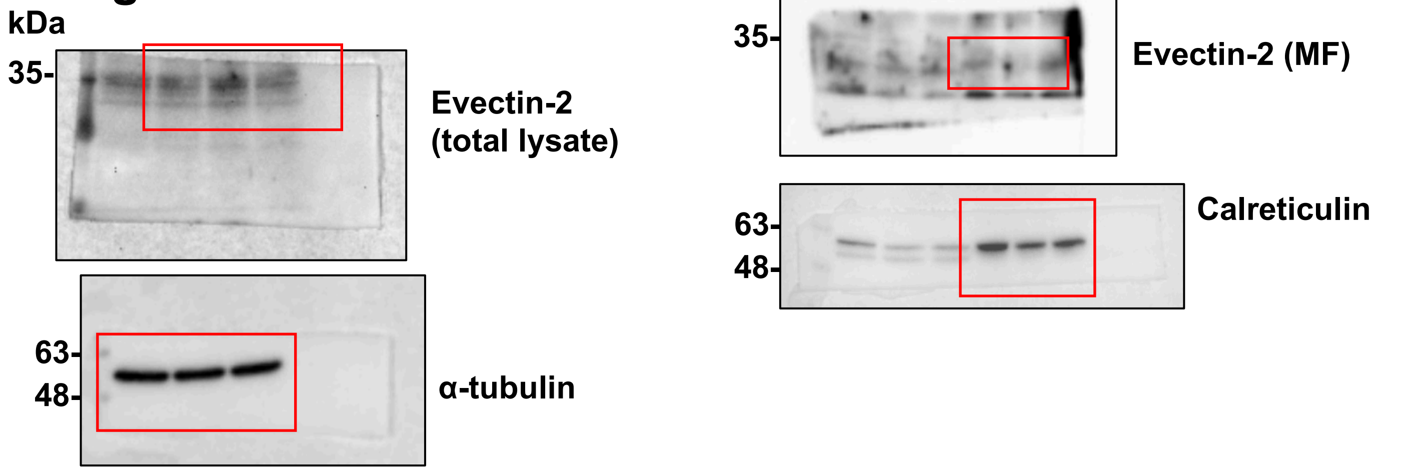


**Supplementary Figure 10 (continued). The original western blot data of Fig. 1~6.**

#### Fig. 3B

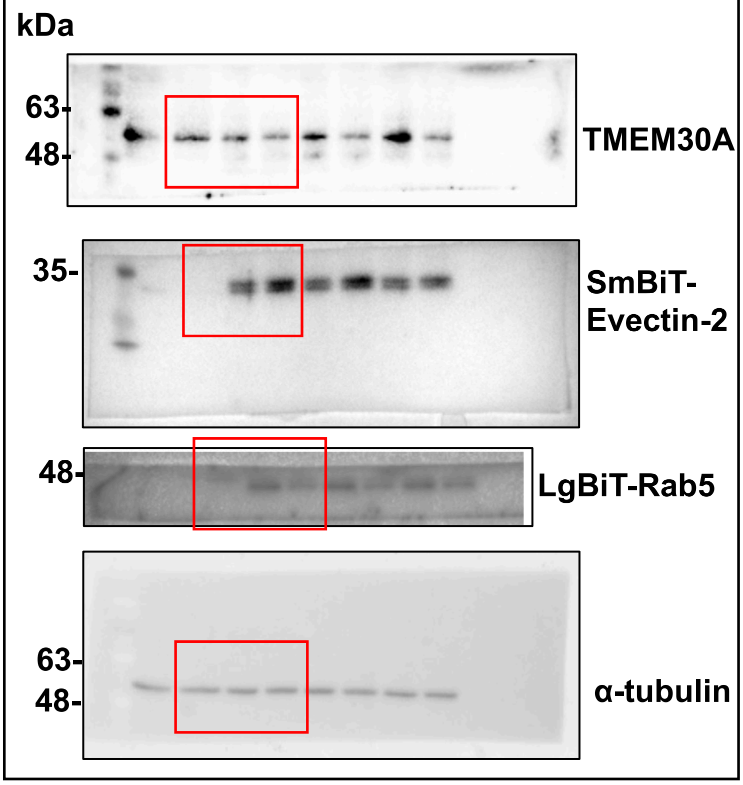


#### Fig. 3D

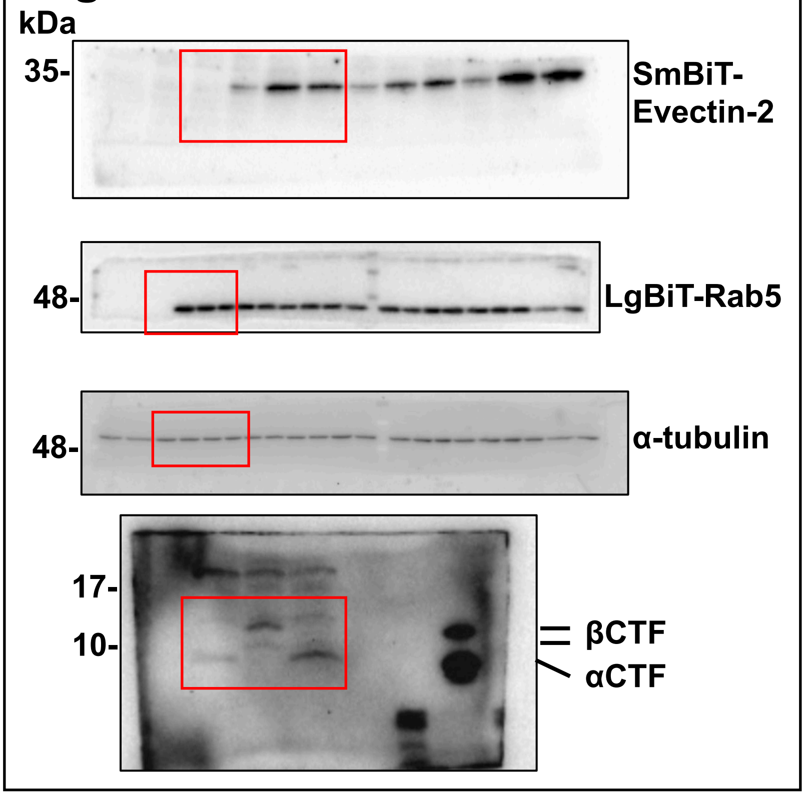


#### Fig. 3F

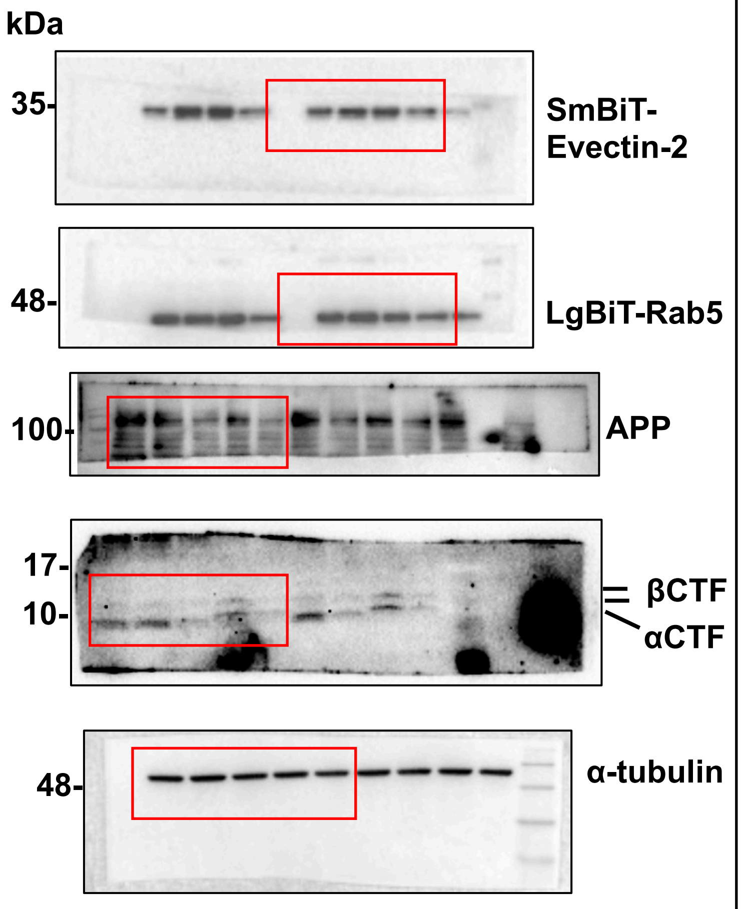


#### Fig. 3H

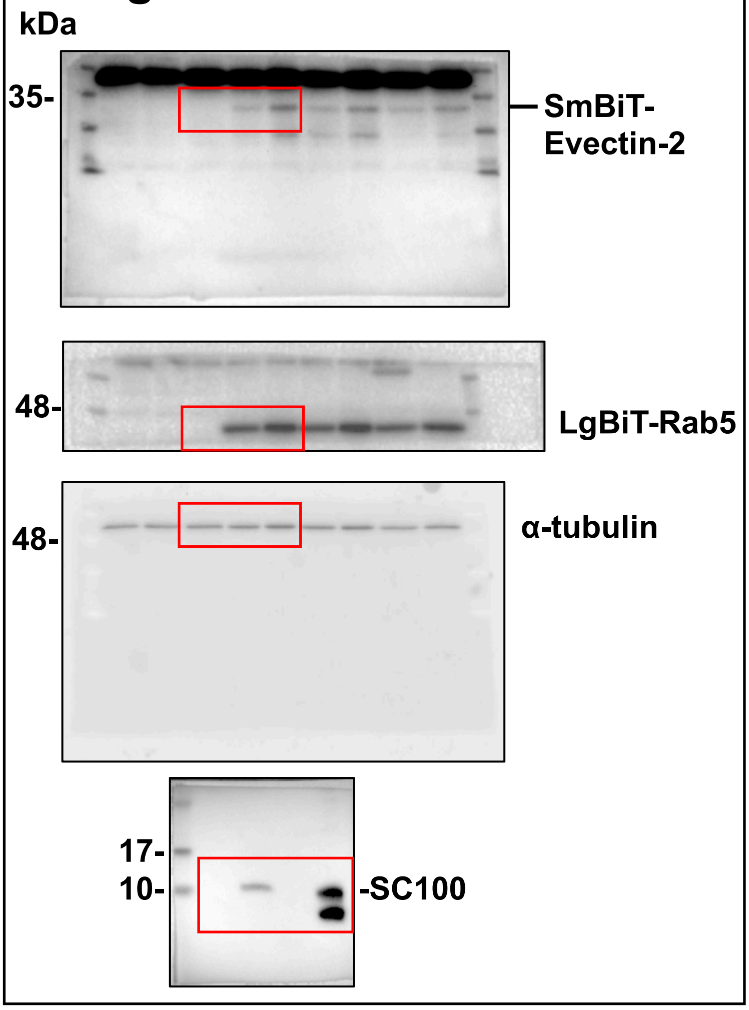


Supplementary Figure 10 (continued). The original western blot data of Fig. 1~6.

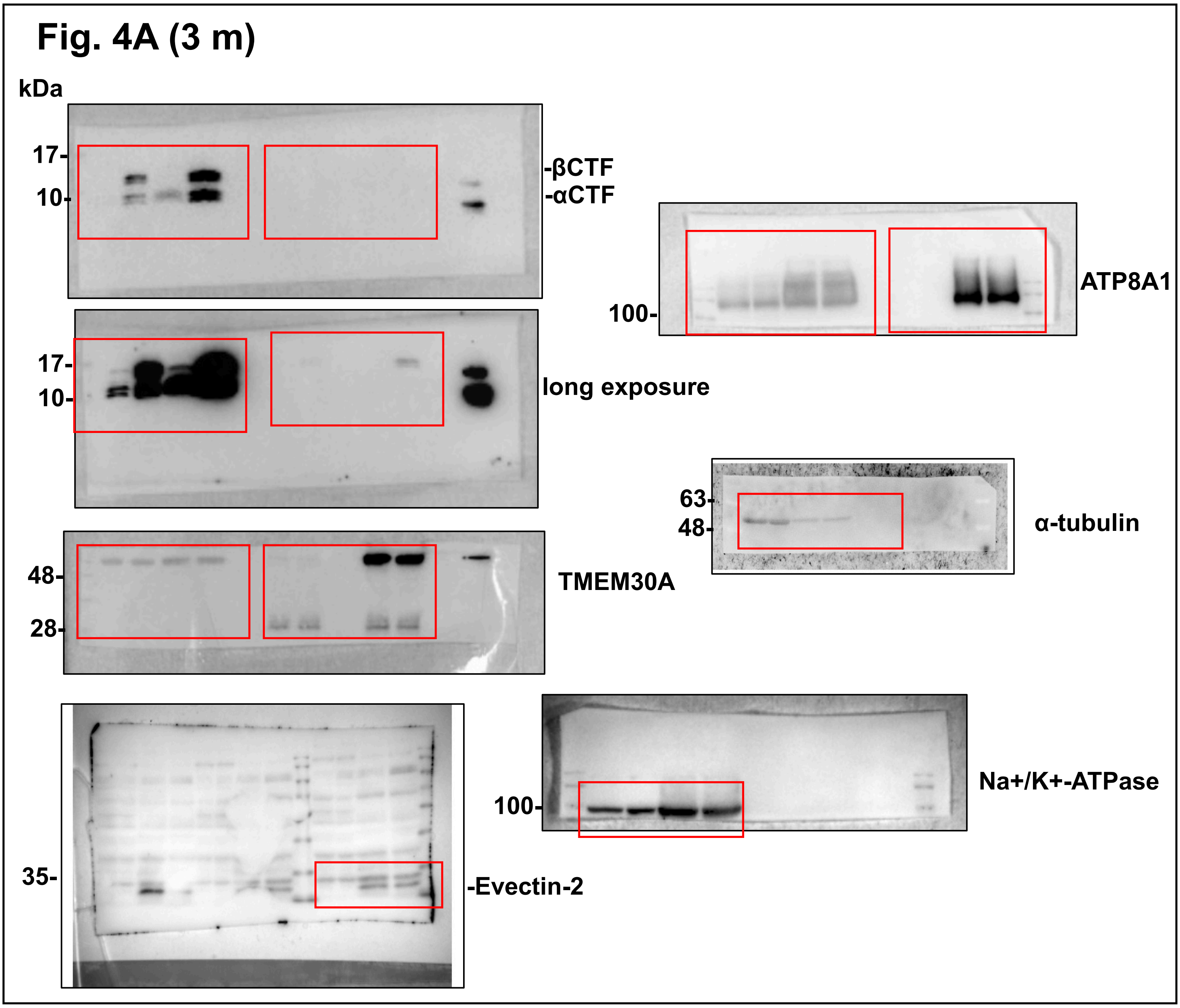


Supplementary Figure 10 (continued). The original western blot data of Fig. 1~6.

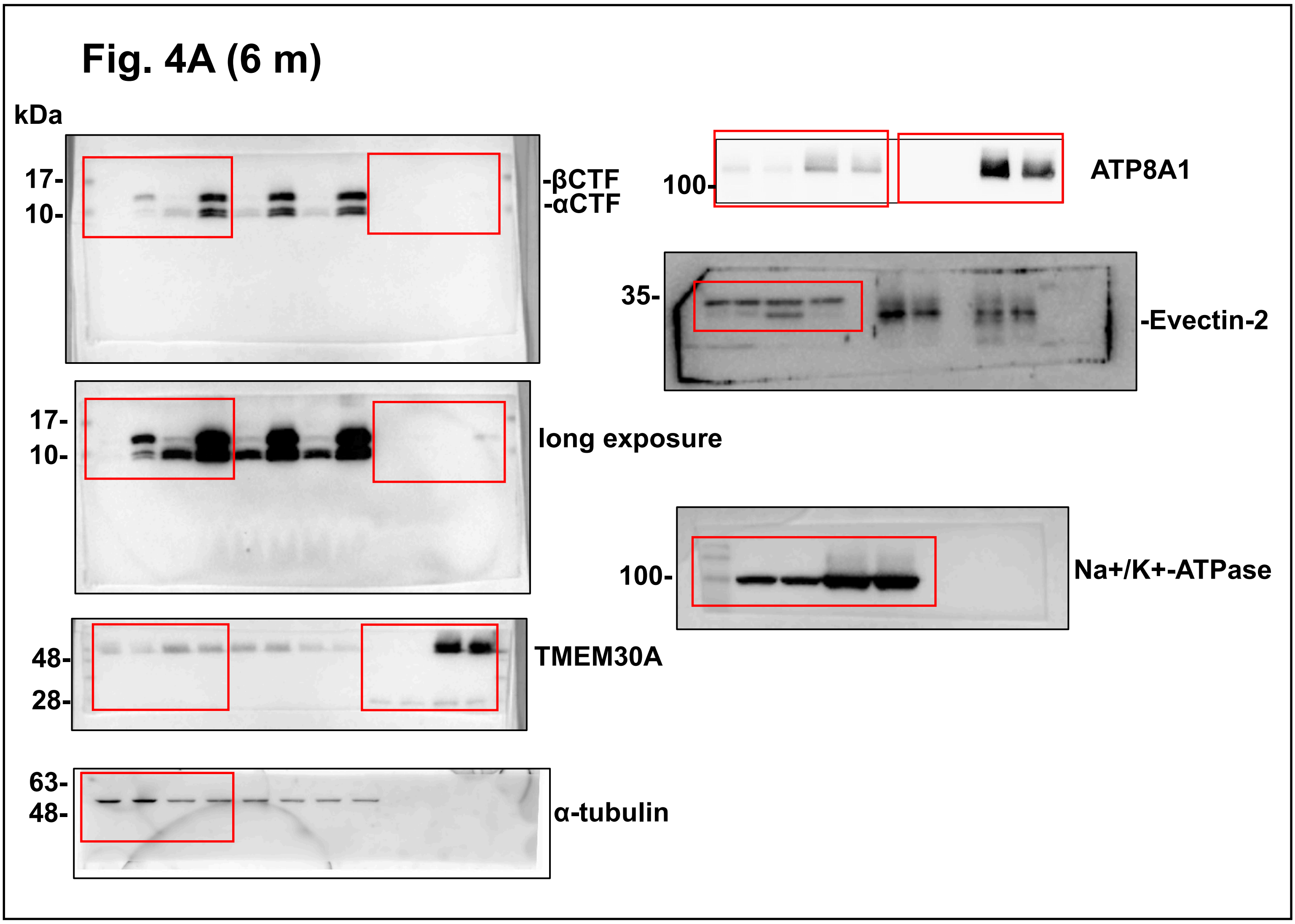


**Supplementary Figure 10 (continued). The original western blot data of Fig. 1~6.**

### Fig. 5B (CBB staining)

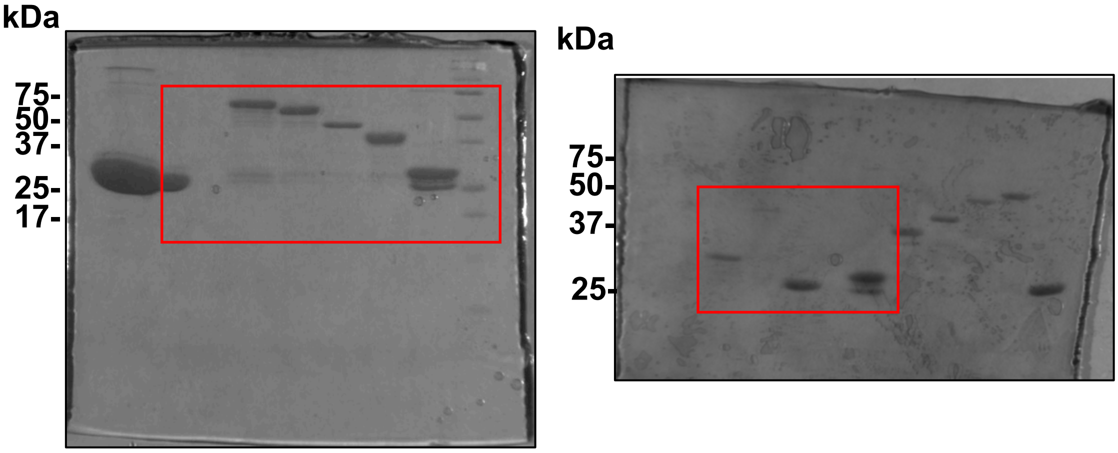


### Fig. 5C

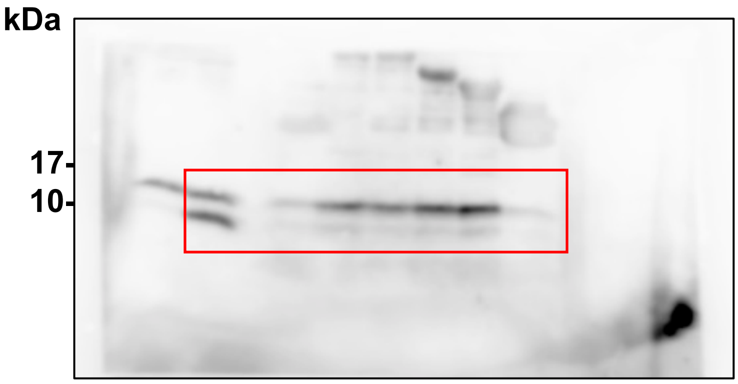


### Fig. 5D

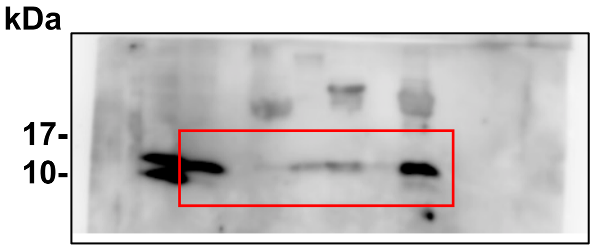


**Supplementary Figure 10. The original western blot data of Fig. 1~6.**

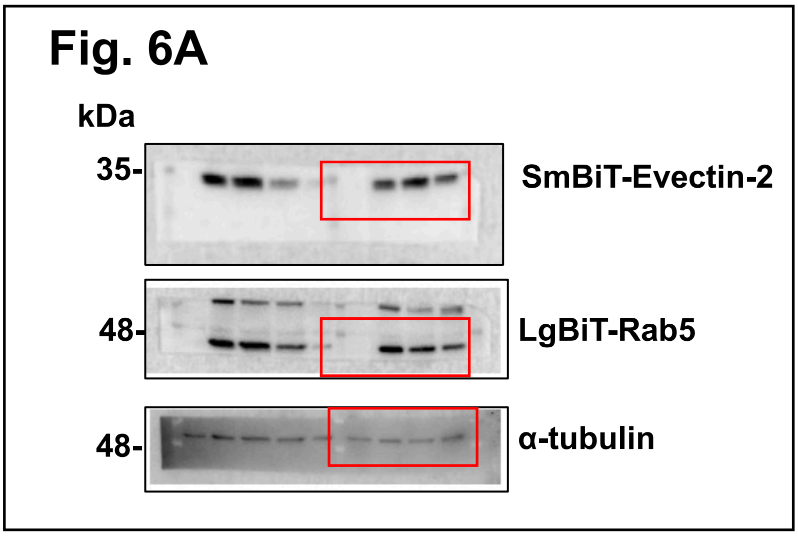


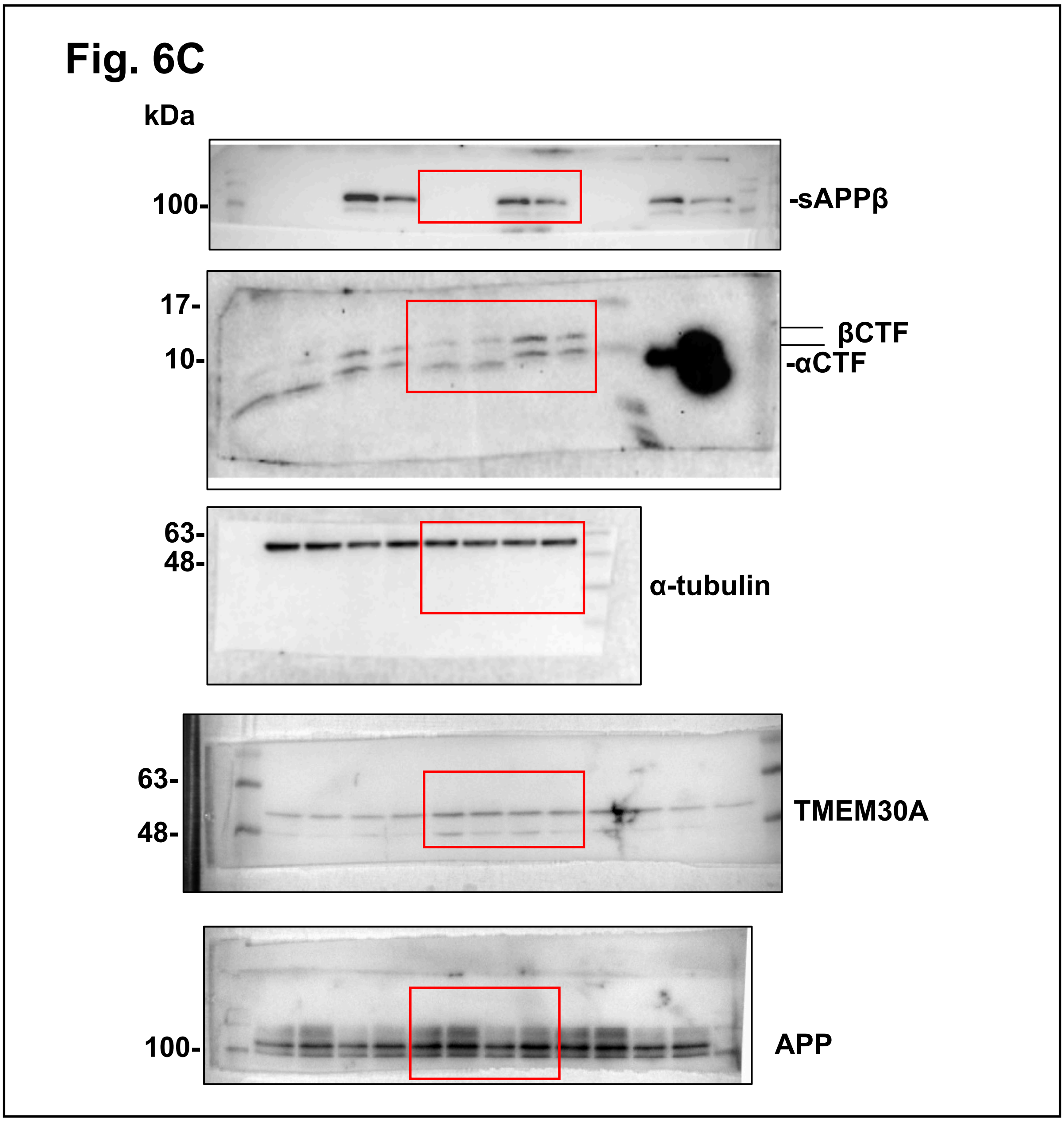


**Supplementary Figure 10. The original western blot data of Fig. 1~6.**
